# Multi-modal image analysis for large scale cancer tissue studies within IMMUcan

**DOI:** 10.1101/2024.12.16.628597

**Authors:** Nils Eling, Julien Dorier, Sylvie Rusakiewicz, Robin Liechti, Preethi Devanand, Michelle Daniel, Jonas Windhager, Bruno Palau Fernandez, Sophie Déglise, Lucie Despland, Abdelkader Benyagoub, Alexander Loboda, Daaf Sandkuijl, Nikesh Parsotam, Henoch S Hong, Marie Morfouace, Nicolas Guex, George Coukos, Bernd Bodenmiller, Stephanie Tissot, Daniel Schulz

**Affiliations:** Department of Quantitative Biomedicine, University of Zurich, Zurich, Switzerland; Institute of Molecular Health Sciences, ETH Zurich, Zurich, Switzerland; Bioinformatics Competence Center, University of Lausanne, 1015 Lausanne, Switzerland; Department of Oncology, Centre Hospitalier Universitaire Vaudois, Lausanne, Switzerland; Ludwig Institute for Cancer Research, Lausanne branch, Lausanne, Switzerland; Immune Landscape Laboratory, Centre Thérapies Expérimentales (CTE), Centre Hospitalier Universitaire Vaudois, Lausanne, Switzerland; Vital-IT group, SIB Swiss Institute of Bioinformatics, Lausanne, Switzerland; Life Science Zurich Graduate School, ETH Zurich and University of Zurich, Zurich, Switzerland; Standard BioTools Canada Inc., Markham, ON, Canada; Merck KGaA, Darmstadt, Germany; EORTC HQ, Avenue E. Mounier 83/11, 1200, Brussels, Belgium

**Author notes:** Shared first author (Nils Eling, Julien Dorier).

## Abstract

**Motivation:** Multiplexed imaging is increasingly used to study tissue architecture in health and disease. To investigate the cancer tumor microenvironment, typically either tissue micro-arrays or small patient cohorts are used to collect and process data. However, studies performed over the course of years, collecting data from thousands of samples are rare and require specialized workflows to ensure sample throughput and reproducibility for data production and processing. Here, we present two such workflows for multiplexed immunofluorescence and imaging mass cytometry of cancer tissues which are applied to a total of roughly 10’000 samples from 2500 patients over six years.

**Summary:** In cancer research, multiplexed imaging allows detailed characterization of the tumor microenvironment (TME) and its link to patient prognosis. The IMMUcan consortium collects multi-modal imaging data from thousands of cancer patients to perform broad molecular and cellular spatial profiling across five cancer indications. Here, we describe two workflows for multiplexed immunofluorescence (mIF) and imaging mass cytometry (IMC) developed within IMMUcan to enable analysis of thousands of cancer tissues. IFQuant supports web-based, user-friendly, and reproducible analysis of mIF data. High sample throughput for IMC is achieved by optimizing experimental protocols and developing a robotic arm for automated slide loading. We provide a resource of 350’000 manually labelled cells across 180 cancer samples to accurately annotate cell phenotypes in IMC. All major cell phenotypes and tissue structures correlate well between mIF and IMC. These pipelines form the basis for multiplexed image analysis within IMMUcan and provide computational tools for the larger community.

## Introduction

The emergence of successful immunotherapy has revolutionized cancer treatment in recent years. However, good prognostic markers for immune response in patients are still lacking and some patients even acquire resistance^1,2^. The TME is composed of tumor cells, immune cells, fibroblasts and endothelial cells, has pivotal anti- and pro-tumorigenic functions^3,4^ and it is commonly accepted that the context of tissue architecture is crucial. Therefore, to reveal the immune cell content of the TME, technologies have been developed to spatially characterize cells in tissue sections^5^. Numerous studies applied multiplexed imaging to investigate the tumor microenvironment and identified signatures of poor or good survival, or predictive of treatment^6–15^. However, efforts to profile thousands of cancer patients with spatially resolved single-cell technologies and molecular data do not exist. To systematically profile the variation of the TME and to characterize biomarkers for diverse treatments, the Integrated iMMUnoprofiling of large adaptive CANcer patient cohorts (IMMUcan) consortium acquires molecular and cellular profiles of over 2500 patients across five cancer types until 2026^16^. By integrating single-cell data obtained from multiplexed imaging technologies with bulk sequencing and whole exome sequencing data and their associated clinical metadata, factors for improved patient stratification or treatment prediction can be unveiled. Within IMMUcan, formalin-fixed and paraffin-embedded (FFPE) tumor samples are processed for molecular and cellular profiling. To study single cells within their spatial tissue context, mass spectrometry based technologies such as IMC^17^ (Standard Biotools) and multiplexed immunofluorescence^18^ (mIF) are established technologies. Both technologies produce complementary read-outs capturing the spatial distribution of dozens of biomolecules, including proteins. While mIF allows the detection of 6-7 fluorescence read-outs across the whole cancer tissue, IMC captures ∼40 proteins in smaller (∼1mm^2^) regions of the tissue. The IMMUcan consortium generates thousands of images using both IMC and mIF to thoroughly profile the TME across different tumor types and thousands of patients. This endeavor raises a number of challenges. First, scalable and reproducible software tools need to be provided to analyze thousands of mIF images with up to millions of cells per image. Second, robust and informative selection of regions of interest (ROI) for IMC^19,20^ needs to be performed. Third, cell phenotypes need to be correctly detected across images, patients and cancer types^21–26^. Finally, a key challenge for a large-scale project such as IMMUcan is to ensure the reproducible processing and analysis of thousands of individual samples over a period of six years.

Here, we present scalable workflows and computational tools to reproducibly analyze multiplexed imaging data of thousands of cancer samples at cellular resolution. We developed the *IFQuant* software for whole-slide semi-automatic quantification of major immune and tumor cell phenotypes in mIF images. The software is embedded within the Knowledge Management System of the IMMUcan project to ensure traceability and reproducibility, and is operated via a web interface for manual curation of images. We further demonstrate automated, robotic sample loading for the IMC Hyperion Imaging System to enable efficient large-scale sample processing. Finally, we show that IF-/metal-antibody co-staining allowed the selection of ROIs for IMC based on whole slide IF images. To facilitate this process with minimum hands-on time, we developed *immucan-roi*, a plugin for the napari image viewer^27^. The acquired IMC data was processed for single-cell extraction and cell phenotyping using a scalable computational workflow and achieved reproducibility across years of sample reception.

Our data analysis workflows were developed independently to optimise processing of either mIF or IMC data. Here, we perform a quantitative comparison of the results of the two workflows and show high correlation and spatial co-localization for all major cell phenotypes between mIF and IMC for samples from five cancer indications. Alongside this we provide a ground truth dataset of IMC images from 179 patients with manually annotated labels of 14 cell types for further methods development. Together, these dedicated, large-scale data acquisition, handling and analysis approaches thus result in accurate implementation of both mIF and IMC to large patient cohorts and ensure high quality and comparability of the data for downstream analysis and characterization of the TME.

## Results

### Highly multiplexed imaging for broad immune profiling of cancer samples

The broad profiling team of the IMMUcan project collects and analyses highly multiplexed imaging (mIF and IMC), whole exome sequencing and RNA sequencing data from over 2500 cancer patients to obtain an in-depth understanding of the tumor immune microenvironment. Patients enrolled in IMMUcan suffer from one of five cancer types: breast cancer (BC), renal cell carcinoma (RCC), head and neck cancer (SCCHN), colorectal carcinoma (CRC), or non-small cell lung cancer (NSCLC) (**Figure 1A, Methods**). The amount of imaging data and sample heterogeneity comes with specific challenges in reproducibility and data integration that require dedicated data acquisition and analysis pipelines.

**Figure 1:**
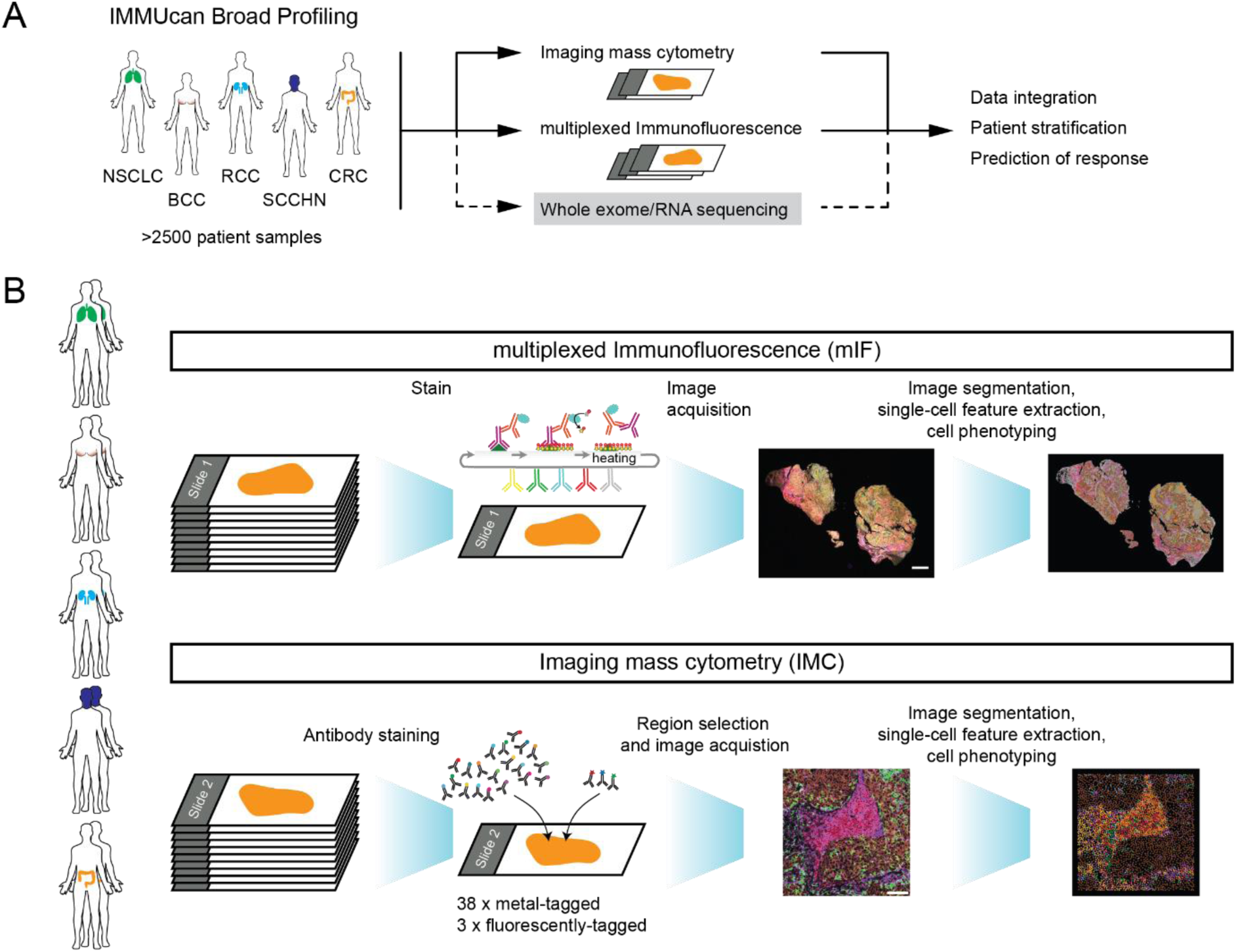
Multiplexed imaging for broad immune profiling of cancer tissues. **A.** The IMMUcan consortium collects and analyzes cancer tissues of more than 2500 patients diagnosed with one of five cancer types: breast cancer (BC), renal cell carcinoma (RCC), head and neck cancer (SCCHN), colorectal carcinoma (CRC), and lung cancer (NSCLC). The broad profiling team of IMMUcan uses IMC, mIF, whole exome sequencing and bulk RNA sequencing, the later two not being presented in this study, to profile the tumor immune microenvironment across all collected cancer samples. Multi-modal data integration will support precise patient stratification and may facilitate the prediction of therapy response. **B.** Ten samples across all five cancer indications were selected to demonstrate the data generation and analysis workflows for IMC and mIF. The first consecutive slide of each patient was stained with 6 antibodies plus DAPI using the PhenoImager™ HT for whole slide fluorescence detection. Image analysis included cell segmentation, feature extraction and cell phenotyping. The next consecutive slide of each patient was stained with 38 metal-tagged antibodies and 3 fluorescent-labeled antibodies. 38 ROIs following general criteria (Methods) were selected across the ten samples for acquisition with IMC. Computational analysis was performed to segment individual cells in images, extract spatially resolved single-cell features and phenotype the cells. Scale bars: 2mm for mIF, 100µm for IMC. Image composites of epithelial cell marker (cytokeratin (mIF), E-cadherin (IMC), brown), CD20 (B cells, red), CD3 (T cells, magenta) and CD163 (macrophages, green) are shown.

We illustrate the mIF and IMC data generation and evaluation pipelines developed within the IMMUcan project using 10 samples that covered each of the five cancer indications (**Figure 1B**). For each patient, we stained the first tissue section for mIF with a 6-plex antibody panel + DAPI using the Tyramide Signal Amplification (TSA) technology^18^ (**Supplementary Table 1, Supplementary Fig 1**) and acquired images using the PhenoImager™ HT. The next, consecutive tissue section of each patient was stained for IMC with 38 metal-labeled and three fluorescent-labeled antibodies (**Supplementary Table 1**). IMC is limited in throughput and requires the definition of regions of interest (ROIs). The fluorescent stain allowed us to perform guided selection of 38 regions of interest (ROIs) across the 10 samples resulting in 13.7 mm^2^ ablated tissue area **(Methods**). The computational workflows for mIF and IMC data analysis are similar and include sample tracking, image processing and segmentation, single-cell feature extraction, and cell phenotyping as well as tissue structure detection (**Figure 1B**).

### IFQuant Experimental and Quantification Workflow

To profile the spatial distribution of major immune phenotypes across whole tissue slides, we optimized a 6-plex panel of antibodies (CD3, CD11c, CD15, CD20, CD163, cytokeratin (CK), **Supplementary Table 1**) supplemented with DAPI for nuclear staining. Using the Ventana autostainer and the TSA OPAL technology^28^ combined with the PhenoImager™ HT (**Figure 2A**, **Methods**), we acquired images from the ten selected patient samples. Since we expected more than 9600 samples to be analyzed (3200 samples from 2500 patients with three mIF panels each) for the whole IMMUcan project, and given that most existing tools operate as standalone applications and struggle with handling such extensive datasets, we designed and developed the IFQuant software with a specific focus on analyzing mIF data from thousands of patient samples in a manner that is streamlined, reproducible, scalable, and user-friendly. IFQuant is a semi-automatic, web-based mIF analysis software (**Figure 2**) producing standardized data output. Within IMMUcan, the IFQuant software is fully integrated in a laboratory information management system (LIMS) which tracks the reception, staining, scanning, analysis and reporting of the samples and thus, reduces the risk of operator induced errors. A stand-alone version of the software is available as a Docker container and a detailed description of the IFQuant analysis workflow can be found in Supplementary Methods 1.

**Figure 2:**
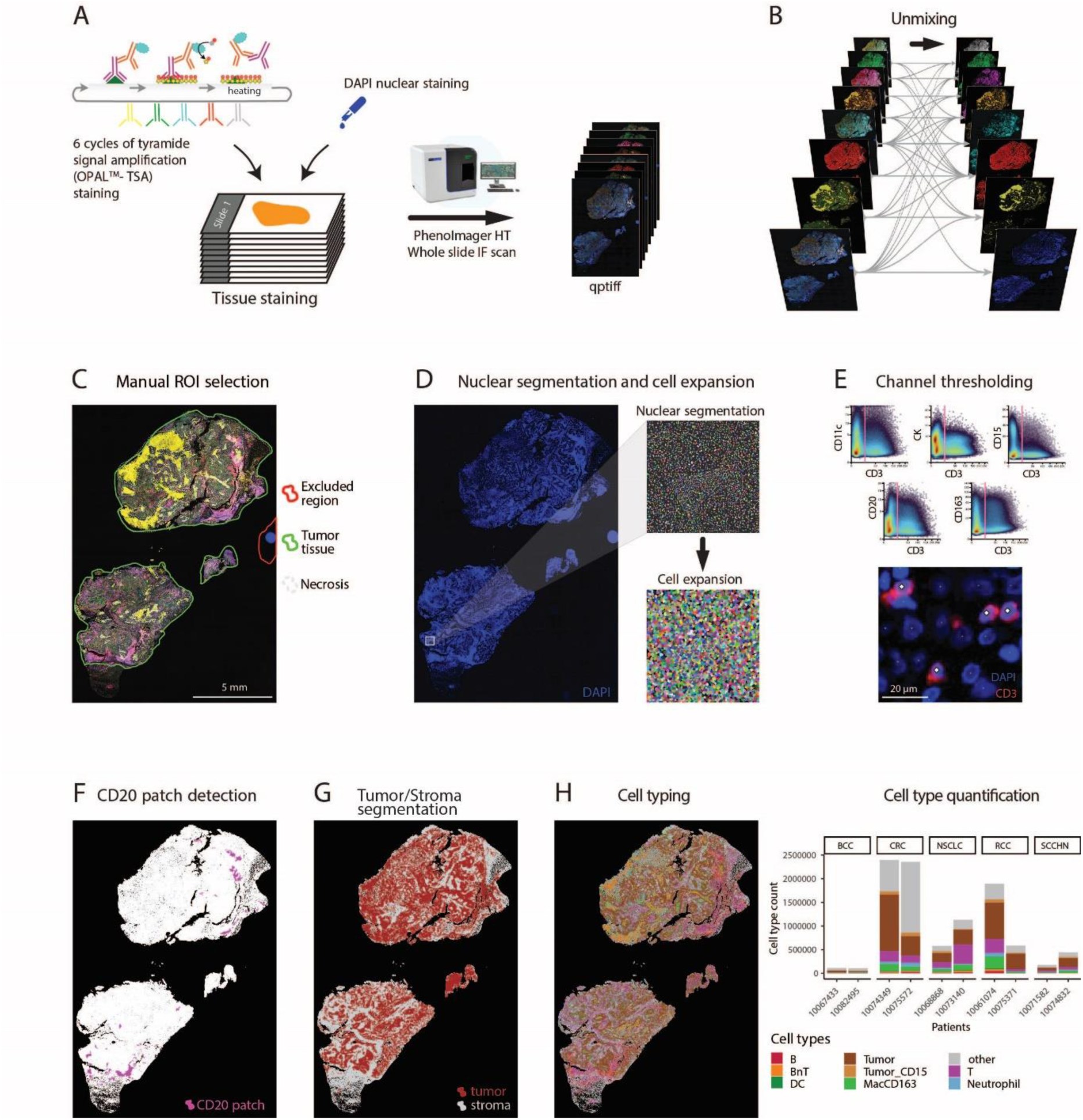
Overview of mIF experimental and IFQuant image analyses workflow. **A**. Tissue staining was performed using 6 cycles of tyramide signal amplification with specific antibodies and DAPI for nuclear staining. Slides were scanned using the PhenoImager™ HT. Quantification workflow: Images were spectrally unmixed (**B**) and annotated (**C**) using IFQuant. **D**. Single cells were defined using DAPI based nuclear segmentation followed by nuclear mask expansion for single cell detection. **E**. Channel-specific thresholding is performed manually to identify marker positive cells. **F**. CD20 patches are detected as proxy for TLS, based on CD20 signal (shown in pink). **G**. Areas of predominant tumor or non-tumor cells are detected and annotated automatically (tumor depicted in red and stroma in grey). **H**. Identified cell types based on marker positivity are shown in the image (left panel) together with a quantification of total cell counts per sample and cell type (right panel).

IFQuant directly accesses QPTIFF image files from the LIMS and performs multi-channel signal unmixing (**Figure 2B**) for downstream analysis (**Methods)**. Subsequently, IFQuant provides an integrated web interface to facilitate manual analysis steps that cannot be automated. The operator can use a free-hand drawing tool to assign manual annotation masks to the images to either filter unwanted regions (e.g., out of focus, bad tissue quality) or label specific tissue regions (e.g., tumor,necrosis, healthy tissue, adipose tissue, **Figure 2C**). Nuclear segmentation is thereafter performed automatically based on the DAPI signal and single cells defined based on an expansion of the nuclear mask (**Figure 2D**). Cellular phenotypes are assigned based on marker expression. Due to sample-to-sample variations in tissue quality and potential minor differences in pretreatment, fluorescence-based marker signal intensity ranges can vary. Therefore, the operator utilizes the IFQuant web interface to refine thresholds for each marker to extract positive and negative cells which can simultaneously be visually inspected. This adjustment is facilitated by a high-resolution image viewer that presents cell overlays and density scatter plots of marker intensities. (**Figure 2E**).

IFQuant automatically detects B cell patches as proxies for tertiary lymphoid structures based on the threshold for CD20 positivity (**Figure 2F**). Furthermore, IFQuant automatically detects areas consisting mostly of tumor cells (CK positive cells) and areas consisting mostly of stromal and immune cells (CK negative cells) and generates a tumor-stroma mask thereof (**Figure 2G**). During post-processing, the mIF marker panel allows annotation of cells as: B cells, BnT cells (cell positive for CD20 and CD3), T cells, dendritic cells (DCs), neutrophils, macrophages (MacCD163), “other” and epithelial tumor cells (**Figure 2H**). The phenotype key, linking cellular phenotype and marker positivity is shown in **Supplementary Table 2**. IFQuant outputs one tabular file containing marker expression, binary marker positivity, the XY location for each cell, and information regarding annotations (e.g., in tumor, necrosis, etc) for downstream computational analysis.

In conclusion, the IFQuant software streamlines the management and analysis of mIF imaging data requiring 10 to 30 minutes of operator time per tissue section (depending on the tissue size and complexity), and delivers a comprehensive and standardized output of spatially resolved single-cell data for subsequent analysis.

### Frozen antibody mixes enable IMC measurement comparison over time

Individual sample stainings can be a source of batch effects. To minimize such effects throughout the IMMUcan project we worked with large antibody mixes sufficient to stain 500 patient samples (**Methods**)^29,30^. For each set of 19 patient samples, we stained and acquired one slide containing two control cell pellets. One cell pellet contained a mix of an epithelial cell line and non-activated PBMCs. The other cell pellet contained epithelial cells cultured with IFN-ɣ, thereby upregulating the expression of PD-L1, and activated PBMCs (**Methods**). To investigate the reproducibility in cell type detection and marker expression over time, the images acquired from control cell pellets over the course of more than 2 years were processed, and single cells were clustered and annotated to identify cell phenotypes (**Figure 3A, Methods**). Epithelial cells were most abundant and separated based on IFN-ɣ treatment while PBMCs separated by cell phenotype and PHA treatment (**Figure 3A, B, Supplementary Figure S2A).** Additionally, we observed a small batch effect based on the antibody staining mix visible on non-activated epithelial cells (**Supplementary Figure S2B, C**). The fraction of detected cell phenotypes in the cell pellets showed a mean coefficient of variation of 15% and 25% for cell phenotypes in activated and non-activated cell pellets, respectively (**Figure 3C, Supplementary Figure 2D**). We compared the expression of relevant markers for cell phenotypes across batches and similarly found a mean coefficient of variation of 16% and 25% for activated and non-activated cell pellet samples, respectively (**Figure 3D**). The variation in staining was also highly correlated with the expression of individual markers, implying that lower signal intensities require larger statistical power for differential detection (**Supplementary Figure S2E**). Overall, these results showed that data acquired over two years is comparable suggesting that cancer images acquired during the IMMUcan project will be comparable.

**Figure 3:**
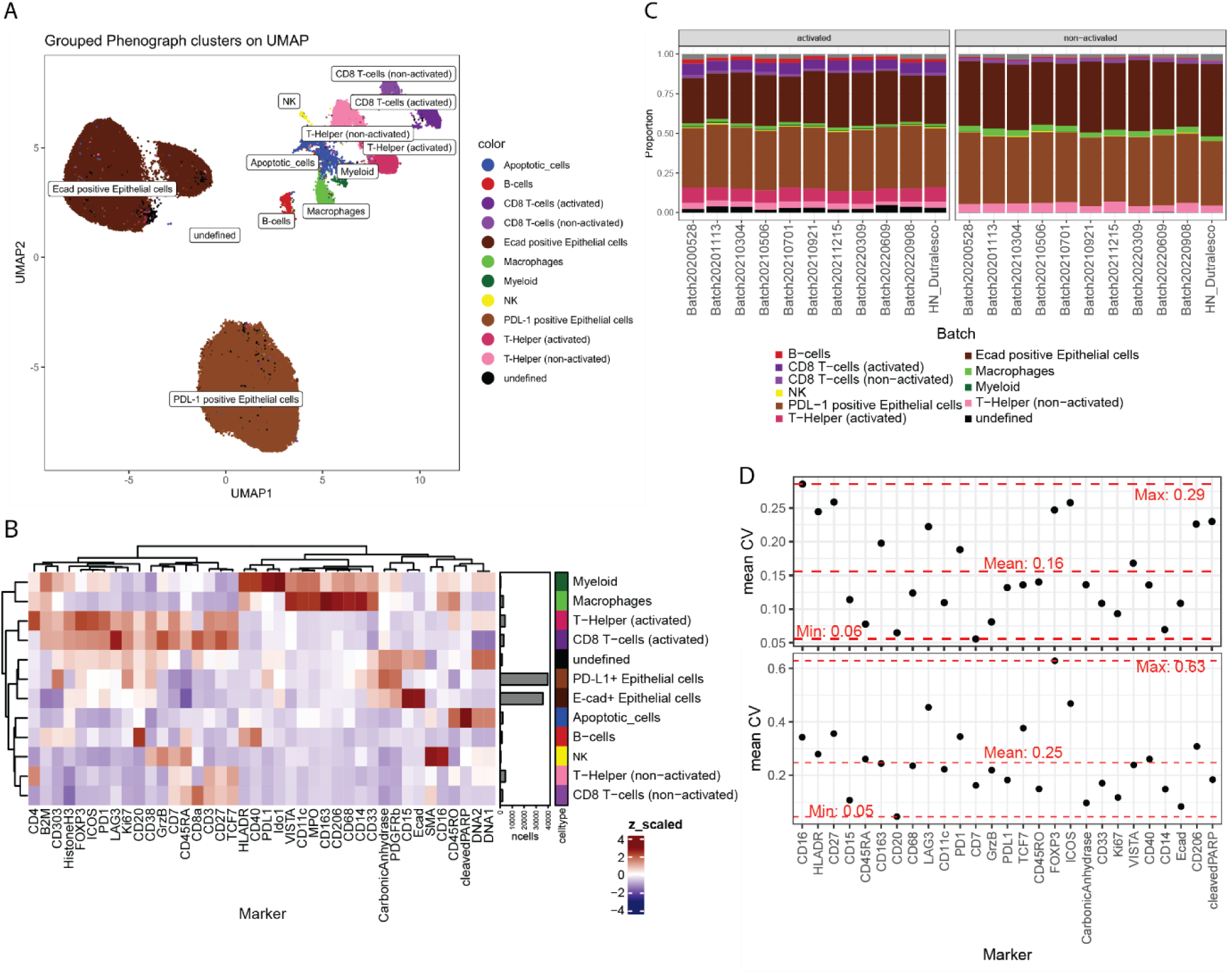
IMC control cell pellets identify variation over time. **A.** UMAP of all cells acquired from sections of cell pellets between May 2020 and September 2022. Cells are colored and labeled according to cell phenotypes. **B.** Heatmap of z-score scaled marker expression averaged per cell phenotype. Barplots show the abundance of each cell phenotype in the data. **C.** Barplot showing the proportions of cell phenotypes of the cell pellets for each sample batch acquired in over two years. Fractions are colored according to cell phenotype. **D.** For each marker expressed in a certain cell phenotype the mean coefficient of variation (CV) over all time points (sample batches) is shown (black points) on top for activated and on the bottom for non-activated cell pellets. The Horizontal red dashed lines indicate the maximum, minimum and mean observed CV across all markers.

### Optimized IMC/IF co-stain for region selection and high throughput of whole slide samples

To stain, measure and analyze thousands of tissue sections using IMC in a reproducible and scalable fashion over six years, we optimized an experimental and computational workflow enabling an efficient throughput for large numbers of samples (**Figure 4**). IMC acquires images at about 1 mm^2^ per hour and therefore requires the selection of regions of interest for multiplexed profiling of whole slide tissues. As current IMC machines only allow the acquisition of bright field overview images (Panoramas) for ROI selection, whole slide immunofluorescence staining and imaging prior to metal-labeled antibody staining have been used in the past for supervised ROI selection^31–33^. However, these multi-step staining protocols are too laborious for the throughput of the IMMUcan project. Therefore, we evaluated the possibility of simultaneously co-staining primary, fluorescent-labeled antibodies with metal-tagged antibodies and detecting the fluorescent signal in a dried tissue format as required for IMC (**Figure 4A**). We selected antibodies against CD45, CD163 and pan-Cytokeratin to identifyimmune cells, macrophages and epithelial cells, respectively. We tested the signal stability of the fluorophores after sample drying and did not observe a decrease in signal intensity within the first 48 hours after tissue drying (**Supplementary Figure 3A, B**). In addition, we optimized for fluorescent-labeled antibody clones against CD45 and CD163 that did not interfere with the metal-labeled antibodies targeting CD45RA, CD45RO and CD163 (**Methods**). Suitable, fluorescent-labeled antibodies were added to the metal-labeled antibody mix for standard staining of IMMUcan samples (Supplementary **Table 1, Figure 4A**).

**Figure 4:**
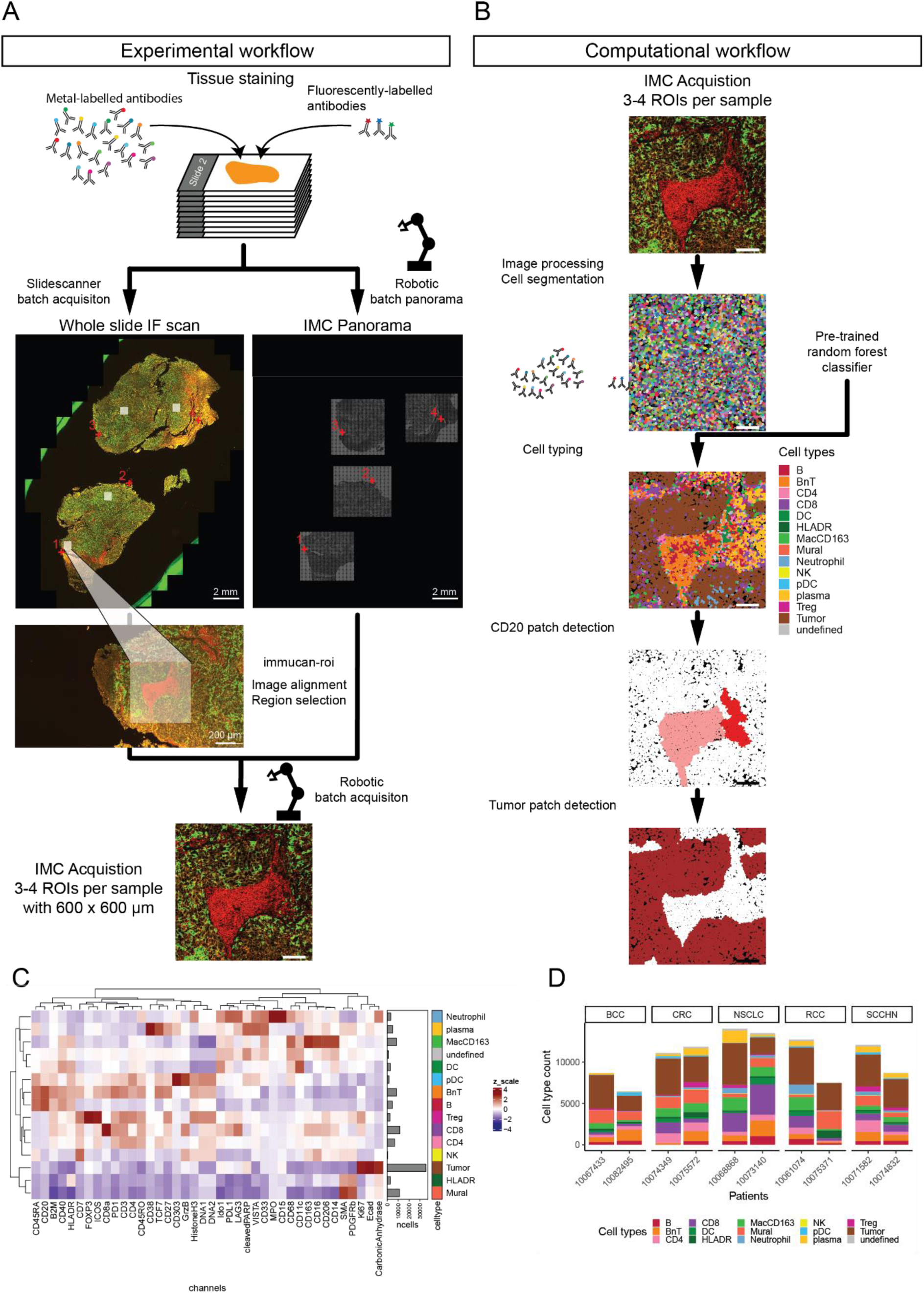
IMC workflow for high throughput acquisition and analysis of whole tissue slides. **A.** Tissue sections are stained using metal-labelled and fluorophore-labelled antibodies. After staining and drying, slides are immediately scanned overnight using a slide scanner and subsequently a slide loader is used to generate panoramas for IMC. The fluorescence whole slide scan and panoramas are then used to align the two modalities. Once aligned, regions of interest are selected on the IF image and automatically transformed into IMC coordinates. The slide loader is used to acquire the selected regions of interest in batches of up to 40 slides with variable number of ROIs per slide. **B.** The computational workflow encompasses pre-processing and single-cell segmentation. Cell phenotyping is performed with a pre-trained classifier and patches of B cells and tumor cells are detected. **C.** Heatmap showing the z-score scaled mean marker expression in cell types classified in the 10 samples used in this study. Barplots indicate the abundance of each cell phenotype in the data. **D**: Barplots showing the cell phenotype counts per sample.

### Automated slide loading and region selection for IMC

The Hyperion Imaging System for IMC supports the measurement of one slide at a time. To increase the throughput, and scale to the measurement of thousands of slides, we collaborated with Standard BioTools to develop a slide loader for batch processing of microscopy slides. The Hyperion Imaging System and its software were customized to allow the installation of a robotic arm with a slide hotel carrying up to 40 slides. After staining, whole slide IF scans were performed for batches of slides using a SlideScanner and subsequently panoramas were generated for the same batch of slides using the slide loader and the Hyperion Imaging System (**Figure 4A, Methods**). To select regions of interest for IMC based on the IF slide scan we developed *the immucan-roi* plugin for the *napari* image viewer^27^ (**Methods**). The plugin reads IF images (in CZI format) and panoramas (in MCD format) for the manual selection of landmarks for alignment. After alignment, the operator selected three ROIs for IMC acquisition based on the 4 color IF image. Since we are interested in imaging immune cells in the TME, ROIs were placed in regions containing tumor and immune cells (**Methods**). One additional ROI was placed on dense regions of CD45^+^ cells (potentially tertiary lymphoid structures), if such regions were visible in the IF image. The *immucan-roi* plugin additionally allows the annotation of the selected ROIs. Based on the previous landmark-based alignment, *immucan-roi* transforms the selected ROI locations into the IMC coordinate system. The automatically exported CSV files containing the ROI locations in the IMC coordinate system can directly be imported into MCD files for IMC acquisition requiring no further hands-on time at the Hyperion Imaging System. Finally, we used the robotic slide loader for IMC batch acquisition of the selected ROIs (**Figure 4A**). The *immucan-roi* plugin is currently the standard tool for selecting ROIs and image alignment across all IMMUcan samples for acquisition using IMC.

### Downstream analysis of IMC images and cell phenotyping

We developed a computational workflow for data analysis that includes image pre-processing, cell segmentation, feature extraction, cell phenotyping and spatial analysis for IMMUcan. For raw data processing, cell segmentation and feature extraction we use the *steinbock* toolkit (**Figure 4B**, **Methods**). Most single-cell analysis steps follow our standard workflow (available at https://bodenmillergroup.github.io/IMCDataAnalysis/)^34^. To robustly define cell phenotypes across shipment batches for the IMMUcan project (**Methods**), we performed random forest-based classification. Therefore, we selected 179 images from individual patients from data acquired over almost three years and manually annotated more than 340’000 cells. Images were selected to represent different sample batches and cancer indications. We trained a random forest classifier on 80% of the annotated cells while providing the cancer indication as covariate and validated the classifier on the remaining 20% of the annotated cells (**Methods**). The classifier detected most cell phenotypes with true positive rate (TPR) greater than 0.8 and a false positive rate (FPR) smaller than 0.03 (**Supplementary Figure 4A**). This classifier is applied to all samples stained within IMMUcan to detect the defined cell phenotypes (**Figure 4B**). Based on the detected cell phenotypes we define CD20 patches and tumor patches (**Figure 4B**, **Methods**). In sum, the developed computational workflow is reproducible, scalable and generates results for around 100 samples, containing 0.5-1 milion cells, in around 10 hours as part of standardized IMMUcan sample processing (with parallel processing on a 32 core, 128 GB RAM Ubuntu system).

### Cell phenotypes and spatial features are comparable between IMC and mIF

The experimental and computational workflows for mIF and IMC newly developed within IMMUcan were designed to ensure high reproducibility and throughput, enabling the acquisition of data from thousands of patients. We next compared the results of these two pipelines, to assess whether cell phenotypes and their spatial relations are comparable between the two technologies. This comparison forms the basis for downstream analysis steps within the IMMUcan project, including patient stratification and biomarker discovery based on cell phenotypes and spatial features.

We analyzed nearly 10 million cells in mIF and 100,000 cells in IMC across the 10 samples processed here. While cell numbers from IMC, which analyses roughly equal regions of tissue, were comparable across samples (6,643 to 14,220 cells per sample), we observed larger variation in cell numbers in mIF ranging from around 100,000 cells in biopsies to more than 2 million cells in large surgical specimens. For mIF, cell phenotyping was performed by manual thresholding of marker intensities and annotation of 8 unique phenotypes (**Figure 2H**). Cell phenotyping in IMC was performed by cell type classification and resulted in 14 unique cell phenotypes (**Figure 4D**). Seven of these phenotypes are matched between both technologies: B cells, BnT cells, T cells, dendritic cells, macrophages, neutrophils and epithelial tumor cells. We label B cells that sit in close spatial proximity to T cells as “BnT cells” as they often show strong marker overlap.

To compare the data from mIF and IMC, we aligned the acquired IMC ROIs with the matched regions on the mIF slides using *napari* (**Methods**), identified cells in matched regions, and compared their properties (e.g., cell phenotype) between the aligned images (**Figure 5A**). Overall, we observed a larger number of segmented cells in IMC compared to mIF (**Supplementary Figure 5A**), possibly due to differences in resolution between the technologies and potential differences in segmentation. This was supported by the observation that cells in IMC on average had a smaller area suggesting over-segmentation in IMC (**Supplementary Figure 5B**).

**Figure 5:**
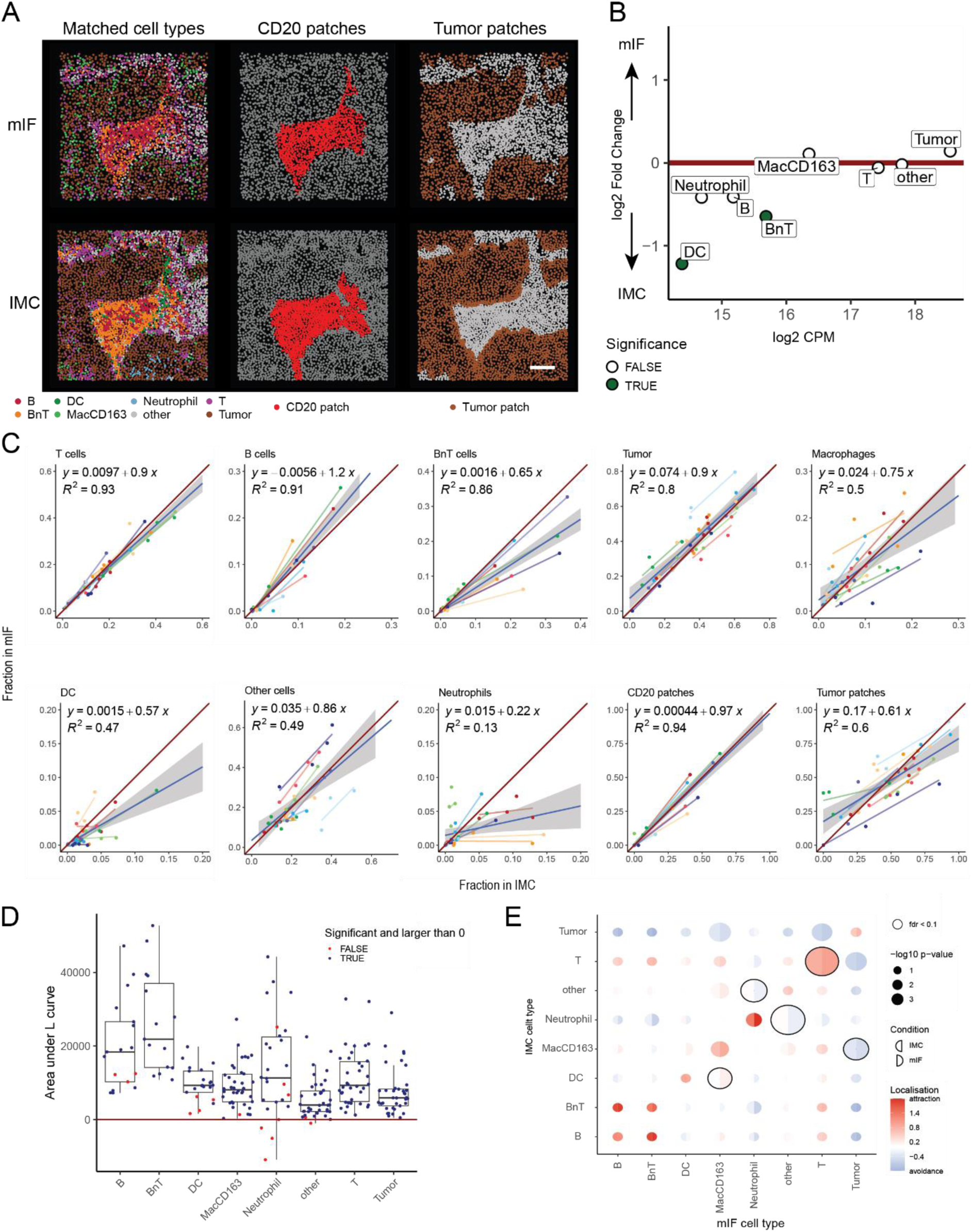
Cell phenotype and spatial feature comparison between mIF and IMC. **A**. Matched image from IMC (top) and mIF (bottom) colored by cell type (left), CD20 patches (middle), tumor-stroma compartment (right). Scale bar: 100 µm. **B.** Differential abundance of cell phenotypes in mIF versus IMC images. **C.** Scatter plots comparing the fractions of each cell phenotype per image in IMC (x axis) against mIF (y axis). **D.** The area between the calculated and theoretical L function for the respective cell phenotype in mIF and IMC. Each dot represents one cell phenotype per IMC-mIF image pair. Blue points indicate cells that significantly co-localize in mIF and IMC compared to random. Red points indicate comparisons with negative area or for which a MAD test did not result in significant co-localisation (**Methods**). **E.** Results showing the spicyR differential co-localization test. Circles are split into results from IMC (left side of circle) and mIF (right side of circle) indicating whether cells are significantly clustered within the modality. Cell phenotypes that show significantly diffent co-localisation between mIF and IMC are indicated by the black lines around the circles.

We next compared the abundance of cell phenotypes between mIF and IMC across all aligned images (**Methods**). We observed no systematic difference between mIF and IMC except for the numbers of detected BnT cells and DCs which were lower in mIF (**Figure 5B, Supplementary Figure 5C**). DCs are the least abundant cell phenotype in the datasets and reliable detection across consecutive sections is therefore the most challenging. Neighboring B and T cells may be easier to separate in mIF images than in IMC due to higher resolution, resulting in lower counts for BnT cells in mIF.

Linear regression analysis was performed to estimate the variance in cell phenotype fractions in IMC that can be explained by cell phenotype fractions in mIF. We observed high similarity in the fractions of T cells, B cells, BnT cells and Tumor cells (R^2^ > 0.8) between mIF and IMC (**Figure 5C, Supplementary Figure 6**). Neutrophils showed the lowest similarity between the methods (**Figure 5C**), which may be explained by the differences in the markers used for neutrophil detection between IMC (MPO^+^CD15^+^ cells) and mIF (CD15^+^ cells). Overall, we conclude that for the most abundant cell phenotypes the abundances in mIF and IMC are comparable.

B cells, and especially B cell patches or TLS are important for immunotherapy prediction^7,35,36^ and thus crucial features to be extracted in images. We compared mIF and IMC in terms of the fraction of cells located within B cell patches per image (**Figure 5C, Supplementary Figure 7A, B**). We first ensured that B cell patches in the images were found in matched locations using a spatial score based on homogeneous L functions^37^ (**Methods**).

A positive area between the observed and the expected L function indicates cells to reside in the same spatial location on matched IMC and mIF images. In addition, we performed a statistical test to assess if the observed L function is significantly different from the expected L function. Even though the methods of B cell patch detection between mIF and IMC differed, the spatial arrangements of the B cells in were similar in IMC and mIF images (**Supplementary Figure 7C, D**). We next assessed whether all matched cell phenotypes reside in similar spatial locations on IMC and mIF images (**Figure 5D**). Overall, for all matched cell phenotypes we observed a positive difference between the observed and expected L function indicating that cells show similar spatial distributions between mIF and IMC. For some images, DCs and neutrophils did not show similar spatial distributions which is in line with the results of the differential abundance test and linear regression analysis. We performed a related spatial analysis testing for differences in cell phenotype/cell phenotype co-localization between IMC and mIF^38^ (**Figure 5E**). Most observed cell phenotype pairs show similar co-localization in IMC and mIF. In line with previous results, DC-macrophage and other-neutrophil co-localization differed between mIF and IMC due to detection variability between the two technologies. In summary, the abundance and spatial distributions of cell phenotypes were similar between IMC and mIF showing the robustness of both analysis pipelines to generate data for downstream analysis.

## Discussion

We present data generation and analysis pipelines for multimodal, high-multiplex imaging of FFPE tumor tissue across research centers and at high throughput. Within the context of the IMMUcan project^16^, these pipelines will be used with three mIF panels and two IMC panels applied to consecutive FFPE tissue sections to evaluate immune infiltration in five different cancer indications. The approaches we have developed will enable reproducible, integrated mIF and IMC analyses in IMMUcan and enable throughput of 3200 samples each for mIF (7500 slides) and IMC (5000 slides) over 6 years.

The importance of immune cells in cancer is beyond doubt. The spatial distribution of immune cells varies across tumor indications or metastatic lesions and has been shown to affect not only disease progression^39–43^ but also to predict responses to treatment^14,35,44–46^. Recent studies have investigated pan-cancer differences of immune cell distributions^47–49^, but no large-scale imaging-based studies exist. IMMUcan uses a pan-cancer immune antibody panel across the 5 different tumor indications to analyse immune cell distributions and to define potential biomarker(s) of disease evolution and response to therapy within and across indications.

Our pipelines enable the integration of two complementary imaging modalities. mIF is capable of imaging whole slide tissue sections, thereby providing an overview of the immune cell distribution within the whole tissue at limited marker resolution. IMC in turn is relatively slow and can therefore only be used to zoom into certain regions of the tissue but measuring 40 markers simultaneously and therefore providing a more detailed characterization. mIF has been broadly applied in different tumor indications^50–52^ but in practice, is not yet available nor used in clinical pathology^53^. Commercial software (Inform - Akoya, HALO – Indica labs, Phenoplex - Visiopharm) has been developed for mIF data analysis^54^, but such products are expensive, have limited throughput and are not designed to handle cohorts of thousands of samples on individual slides. Thus, for the purpose of IMMUcan, we developed IFQuant and integrated it into the laboratory management system. This creates a seamless sample processing workflow including sample tracking, monitoring of the staining and scanning procedures, semi-automatic mIF analysis and batch download of quantification results for downstream computational analysis (**Methods**, **Supplementary methods**). IFQuant allows user-friendly computational analysis of mIF images by segmenting individual cells, extracting spatially resolved single-cell features and single-cell phenotypes.

Previous IMC studies of tumor samples in patient cohorts have mostly relied on small sample numbers or have made use of tissue micro-arrays (TMAs) to study large cohorts^10,12,14,15,41,55^. TMAs are often designed by clinicians and pathologists such that the selected tissue regions are of clinical or biological interest. Two studies have investigated the relationship between tissue sampling and the power to detect cell types and spatial structures^19,20^. These studies found that the probability of detecting a certain feature (cell type or spatial structure) depends on the abundance of the feature, the size of the spatial structure and the sampled tissue area.

For IMMUcan we measure 1 mm^2^ per sample with IMC and distribute it among 3 ROIs of 600x600 µm. In the context of these two previous studies, using this approach will likely not permit the detection of rare events. However, the ROI size allowed us to capture structures such as tumor-stroma interfaces or TLS.

To enable biologically-informed selection of ROIs in a high throughput fashion, we developed an IF/IMC co-staining protocol in which we used fluorescent-labeled primary antibodies to identify epithelial cells, immune cells and macrophages across whole tissue sections, thus drastically reducing hands-on time per slide while maintaining good fluorescence readout for ROI selection. The use of fluorescent-labeled primary antibodies comes at the cost of lower IMC signal intensity and a potential competition between IF and IMC antibodies targeting the same epitope. As an example, we could not identify a well performing fluorescent-labeled CD3 antibody that did not interfere with the signal intensity of the metal-labeled CD3 antibody. To make the region selection more efficient we developed the *immucan-roi* napari plugin for landmark-based alignment of the IF images and IMC panoramas and for the selection and annotation of ROIs. Furthermore, the tool directly outputs CSV files which can immediately be used to acquire IMC data.

The most critical feature enabling the throughput required for this project was the slide loader attached to the laser ablation unit of the Hyperion Imaging System. We collaborated with Standard BioTools to develop a robotic arm for batch acquisition of brightfield panoramas or IMC images of up to 40 slides, enabling continuous data generation.

Computational analysis included image segmentation, single-cell feature extraction and cell phenotyping. While cell phenotypes in mIF were defined based on marker positivity, we trained a random forest classifier on more than 300,000 manually labelled cells across 180 images and 5 cancer indications. We provide this resource to enable future developments of improved cell phenotype classification strategies for multiplexed imaging data.

We identified a set of seven cell phenotypes that matched between mIF and IMC. The overall abundance and spatial location of those cell phenotypes showed high similarity between IMC and mIF, even though the computational approaches for cell phenotyping differed. This comparison ensures that all major cell phenotypes are correctly detected in mIF, therefore enabling the stratification of inflamed or immune excluded tissues. On the other hand, correctly identifying major cell phenotypes in IMC facilitates a precise sub-classification of immune cells such as exhausted T cells or sub-types and states of myeloid cell. We observed discrepancies in the detection of rare cell types, namely DCs and neutrophils, between mIF and IMC. This difference can arise due to IMC data containing more markers for accurate cell phenotype detection. As such, the combination of CD15 and MPO allowed precise detection of neutrophils in IMC while CD15 alone in mIF lead to rare misclassification of neutrophils as tumor cells. One limitation of the presented comparison is the size of region across which we compare cell phenotypes. Due to restricted ROI size in IMC, we performed cell phenotype comparisons across roughly 1mm^2^ tissue regions per patient. While this allowed us to identify similarity in the local distribution of cell phenotypes and smaller tissue structures, we are not able to confirm that cell phenoptype distributions match between IMC and mIF across the whole tissue.

In conclusion, we present data generation and processing pipelines for multiplexed and multimodal imaging of tumor samples in large patient cohorts, in the context of the IMMUcan consortium. We showed that our pipelines operate reproducibly and at high-throughput built on open-source software, and yield consistent cell phenotypes and spatial patterns across the two technologies. These methods will be used within IMMUcan to investigate biological and clinical features of patient groups across indications. Our findings should benefit other large scale tissue imaging and analysis projects and both our tools and single-cell resource of 340,000 phenotypically annotated cells are publicly available for this purpose.

### Limitations of the study

For mIF, IFQuant enabled us to process all the samples. However, the process is still time-consuming and tissue regions have to be assigned manually (e.g. necrosis). While a working solution was needed at project start, in principle we could now use all the annotated data to train deep learning models for automated region annotation and potentially also for single cell phenotyping for the future. For IMC, the selection of ROIs for each patient is likely the most critical part and was performed manually for multiple reasons. First, the criteria for algorithmic ROI selection were not known at the start of this discovery project and are not known by now either. However, our data may help addressing this question in the future. Second, tumor tissue often has adjacent normal tissue which cannot be distinguished purely based on pan-cytokeratin expression, making automated, purely marker-based region selection difficult and error prone. Lastly, manual selection of ROIs also ensures quality control as each tissue is checked during the process. On the downside, manual region selection is time consuming, subjective and will always contain a certain bias. We tried to minimize this bias using selection guidelines as described in the methods.

## Acknowledgements

The IMMUcan project has received funding from the Innovative Medicines Initiative 2 Joint Undertaking under grant agreement No 821558. This Joint Undertaking receives support from the European Union’s Horizon 2020 research and innovation program and EFPIA https://IMI.europa.eu. The SPECTA Platform is supported by Alliance Healthcare. Alliance Healthcare will become Cencora. We thank Natalie de Souza for critically reading the manuscript. We thank Alex Berlyand for mechanical design of slide loader with robotic arm. We thank Nikolai Alexandrov and Olga Loboda for developing a slide loader extension of CyTOF software and its UI for this project.

## Author contributions

DS, NE, SR, ST and RL designed the study. JD and RL developed IFQuant with input from ST and SR. SR, PD, LD and AB performed mIF experiments. JD, SR, PD, LD and AB analyzed the mIF data. AL, D Sandkuijl, NP developed robotic slide loader with input from DS. MD, SD and DS performed IMC experiments. NE, DS and BP performed IMC data analysis. NE, JD and DS performed mIF and IMC comparison. JW developed the *immucan-roi* napari plugin. NG, GC and BB gave input during project progression. NE and DS wrote manuscript with input from all authors.

## STAR Methods

### Patient samples

Clinico-pathological data and samples were collected according to the EORTC-SPECTA protocol (NCT02834884), open in 20 European countries and 126 hospitals. All patients provided written informed consent at the time of sample collection for molecular and cellular analysis. The IMMUcan project was approved by several ethical committees; see below for the main recruiting sites: Commissie voor Medische Ethiek ZNA, Belgium on 10/03/2021; Comite de protection des personnes “Nord-Ouest I”, France on 02/12/2019; Ethik-Kommission der Medizinischen Universität Wien, Austria on 15/04/2020; Cyprus National Bioethics Committee (CNBC), Cyprus on 29/06/2023; Comissäo de Ética CHUPorto / ICBAS, Portugal on 03/11/2021; CES do IPO Porto, Portugal on 11/03/2021. Patient sample 10074349/IMMU-CRC1-0410 was a distant metastatis. Patient material in IMMUcan is distributed in the form of FFPE blocks, which are cut into individual 4 µm thick sections (**Supplementary Figure 1**). Individual cuts are mounted on microscopy slides, which are sent to the individual partners in roughly 4 shipment batches per year. Slides 1, 4 and 5 are stained with three panels for mIF and slides 2 and 3 are stained with two panels for IMC. In this study, cell phenotype distributions were compared between mIF (slide 1) and IMC (slide 2) for matched tissue regions.

### Generation of positive cytoblock controls

Cytoblocks controls combined PBMCs and established SW480 colon cancer cell line. PBMCs were isolated from patient blood by density gradient using Ficoll-Plaque-Plus (GE Healthcare) and immediately cryopreserved in 90% HS (Human Serum) and 10% DMSO (dimethyl sulfoxide) in liquid nitrogen. SW480 cells were thawed and cultured for 2 weeks in two T75 Flask according to ATCC recommendation to obtain tumor cells in exponential growth phase. IFNg (Roche, #11040596001) was added at 200ng/mL to one SW40 Flask for 24h to induce PDL1 expression. In parallel, PBMCs were thawed and seeded in RPMI 1640 with 8% filtered human serum plus 1% penicillin/streptomycin at a concentration of 1x106 cells/mL with (activated) or without (non-activated) 500ng/mL of PHA (Thermofisher #R30852801) incubated for 48h at 37°C and 5% CO2. SW40 tumor cells were harvested from the two T75 Flasks and mixed with either activated or non-activated PBMC (1:1) to a final concentration of 5x107 cells and centrifuged for 5 min at 400xg. Dried cell pellets were resuspended and fixed with 20X Volume of Shandon Formal Fixx solution (Fisher Scientific #7401150) and prepared using the cytoblock kit (Thermofisher #7401150) following manufacturer recommendation. Briefly, 10x106 cells were mixed with 1 drop of reagent 2,1 drop of reagent 1 plus 2 drops of Hematoxylin in Harris (Biosystems AG # 41-1011-00) to get a semi-solid mass. The fixed cell mixture was enclosed in a biopsy bag (Leica # 3801085) and in a universal cassette with foam pads (Biosystems AG # 81-0023-00 & # 81-0241-00) for inclusion using the biopsy program from the Vaccum Infiltration Processor system (Sakura tissue embedding console system). Paraffin blocks of the tissue mixtures were then generated (Tissue-Tek TEC 5) for downstream sectioning.

### mIF staining procedures

An mIF antibody panel was designed to detect immune cell populations. It contains antibodies against CD15: phenotypic marker of neutrophils, CK: tumor marker, CD163: phenotypic marker of macrophages, CD11c: phenotypic marker of dendritic cells, CD20: phenotypic marker of B cells and CD3: phenotypic marker of T lymphocytes. Antibody references and staining conditions are available in **Supplementary Table 1**.

Multiplexed staining was performed on tissue sections of 4 µm thickness on automated Ventana Discovery Ultra staining module (ROCHE). Slides were placed on the staining module for deparaffinization, epitope retrieval (64 minutes at 98°C) and endogenous peroxidase quenching (Discovery Inhibitor, 8 minutes, Ventana). Each round of staining included non-specific site blocking (Discovery Goat IgG and Discovery Inhibitor, Ventana), primary antibody incubation and secondary HRP-labeled antibody incubation for 16 minutes with Discovery OmniMap anti-rabbit HRP (Ventana, # 760-4311) or anti-mouse HRP (Ventana, #760-4310). Covalent dye labeling was then performed using the OPAL^TM^ reactive fluorophore detection (Akoya Biosciences, Marlborough, MS, USA) for 12 min followed by subsequent heat denaturation of the antibodies for a next round of staining. The mIF stained slides were scanned using the PhenoImager™ HT (AKOYA) with the MOTiF^TM^ mode, allowing whole slide multispectral image acquisition. The scanner outputs a multi-channel 8-bit image in QPTIFF format.

### Antibody conjugation for IF-IMC co-stain

Antibodies for IMC were conjugated following the manufacturers protocol. After conjugation the antibodies were stored at the highest possible concentrations but maximally 500 µg/ml in tris-based stabilizing solution (Candor Biosciences) at 4°C.

For IMC-IF co-staining, Alexa-Fluor 488 conjugated antibodies against pan-cytokeratin (Thermo Fisher Scientific # 53-9003-82), DyLight 550 conjugated antibodies against CD45 (Novus Biotechnologicals #NBP2-34528R), and Alexa Fluor 647 conjugated antibodies against CD163 (Abcam #ab218294) were purchased and stored at 4°C.

### IMC antibody panel generation and storage

A large antibody staining mix sufficient for the staining of roughly 500 slides was prepared. The required amounts of metal-labeled antibodies were mixed and diluted with staining buffer (TBS, pH 7.6, 3% BSA, 0.1% Tween) to a predefined concentration (**Supplementary Table 1**). The mix was then aliquoted into differently sized aliquots and the aliquots stored at −80°C until further use.

### IMC staining

A detailed protocol can be found here (https://www.protocols.io/view/imaging-mass-cytometry-antibody-staining-5qpvo5w2dl4o/v1). Briefly, FFPE slides were removed from – 80°C storage and kept at room temperature for at least 10 minutes to equilibrate to room temperature, deparaffinized and rehydrated and stored in TBS (pH 7.6). To reduce the required amount of antibody solution slides were gently tapped sideways and immediately circled with a PAP-Pen to provide a hydrophobic barrier. Slides were then treated with blocking buffer (TBS with 3% BSA and 0.1% Tween) for 1 h at room temperature. Antibody staining mix aliquots were removed from −80°C and thawed on ice. Fluorescent-labelled antibodies were added to the mix of metal-labelled antibodies (**Supplementary Table 1)** and gently mixed. Blocking solution was removed from the slides and roughly 30-80 µl of staining mix was added to each slide depending on the size of the tissue. Slides were incubated at 4°C overnight in a wet chamber. The next day slides were washed in TBS and roughly 100-200 µl of a mix of Iridium intercalator (1 µM) and Hoechst solution (2 µg/mL) were added to each slide. Slides were incubated for 5 min in a wet chamber at room temperature and washed with TBS. Slides were then briefly dipped in ddH_2_O and then dried under air flow.

### Slide-scanning for IF-IMC co-staining and panorama generation

The dried slides after staining were scanned within 48 hours with a Zeiss AxioScan with 10x magnification and filters for DAPI, 488, 555 and 647 using the ZEN software.

Subsequently, brightfield scans (Panoramas) were generated of the same slides using the Hyperion+ Imaging System (Standard BioTools) with a robotic arm (Meca500, Mecadmic) for automatic slide loading. The system is a prototype, was developed with Standard BioTools (former Fluidigm) and required changes to the hardware and software of the Hyperion+ Imaging System. Briefly, after alignment the robotic arm could insert single microscopy slides from a slide hotel into the ablation chamber and place them back to the hotel. The hotel had a maximum capacity of 40 slides. For batch mode, either panoramas or acquisitions had to be defined for each slide of the hotel and could then be run in batch mode for up to 40 slides consecutively.

### IMC region selection using the immucan-roi plugin for napari

Panoramas (stored in the MCD files of each slide) and immunofluorescence whole slide scans (stored in CZI format) were used as input for the open source *immucan-roi* plugin for the *napari* image viewer^27^. For each matched pair of CZI and MCD file images a set of minimally 4 landmark points were selected on both images to identify corresponding points. Due to fluorescence-based and brightfield-based image acquisition being performed on the same tissue slide, even single cells could easily be visually matched for landmark selection. Once 4 matched landmark points have been identified an affine transformation model is applied to align the images.

After image alignment the tool was used to select ROIs on the fluorescence image. Colors and saturations were adapted as needed and regions selected based on the following general criteria: 1) 30-70% tumor content. To fulfill this requirement, regions were mostly placed at interfaces between tumor, stroma and immune cells. 2) Exclusion of necrotic regions based on DAPI stain. 3) If samples were resections, then one ROI was placed on the global front of the tumor and the other ones in the core of the tumor. 4) If CD45 dense regions (potentially TLS) were visible in the whole slide scan one additional ROI was placed on one such regions inside the tumor area if possible. All regions were selected to be of 600x600 µm size except for the 4th additional ROI or if the amount of tissue material prohibited the placement of multiple ROIs. A CSV file was then automatically exported for all regions of interest in IMC coordinate system space.

### IMC data acquisition

For each slide, the ROIs to be acquired with the Hyperion+ Imaging System, defined using the *immucan-roi* tool, were imported into the corresponding MCD files (created during panorama generation) and saved therein. Identical to the batch panorama mode, a batch acquisition mode was incorporated into the CyTOF software which used MCD files and a linking CSV file as input. Using the robotic slide loader, batches of typically 20 slides were acquired for IMMUcan. The ten samples described in this paper were selected from a range of samples acquired over time. Samples were acquired at 400 Hz with 1 µm resolution.

### IF data analysis

mIF image analysis is performed with the IFQuant software. The detailed methodology as well as the list of tools and packages integrated in IFQuant are described in **Supplementary Methods 1**. Briefly, the following steps are performed:

1. **Unmixing:** Multiplexed immunofluorescence images contain 8 channels (one channel per fluorophore spectral band and one channel for the autofluorescence). Due to the overlapping emission spectra of the fluorophores, the signal of each channel is a mix of signals from all fluorophores and from autofluorescence. With the help of a library of single stained images, IFQuant can subtract the background signal as well as the estimated contribution of the other fluorophores to the signal of each channel. This process creates a multiband pyramidal unmixed TIFF file readable by the IIPImage image server, used only for visualization. All subsequent operations are performed on images that are unmixed on the fly.
2. **Nuclei segmentation:** Nuclei segmentation is done on the unmixed image channel with DAPI nuclear staining by applying an adaptive thresholding strategy, followed by the watershed algorithm applied on the distance map to label individual nuclei.
3. **Cell segmentation:** In the absence of a cell membrane staining, the cell regions are approximated by simultaneously extending each nucleus region by up to 5 μm or until touching a neighboring nucleus region (Voronoi based segmentation^56^). In addition to nucleus and cell region, we also define the cytoplasm region as the set difference of cell region and nucleus region.
4. **Per cell fluorescence quantification:** For each cell, several summary statistics of pixel intensities are computed to quantify the signal of each marker in the whole cell, the cytoplasm and the nucleus regions.
5. **Cell phenotype assignment:** IFQuant uses a simple thresholding approach to classify each cell as being positive or negative for each marker. The type of summary statistics used to quantify a marker depends on the marker location and distribution within the cell (some markers **including** CD3 are surrounding the nucleus whereas others **including** CK have a more diffuse location in the cytoplasm). For each marker a score, defined as a combination of region (nucleus, cytoplasm or cell) and a summary statistic over this region, is empirically chosen. A threshold is manually chosen for each marker and each cell is classified as positive or negative for this marker depending on whether the score is above or below the threshold. To help with the choice of threshold, a web application allows visualizing full resolution unmixed images and dynamically highlighting cells positive for the selected marker, as well as to display thresholds on scatter plots of marker scores for all pairs of markers. A phenotype is finally assigned to each cell using the phenotype key (**Supplementary Table 2**), which maps each combination of marker positivity status to a phenotype. Of note, cells can for example be positive for CD163 and CD3 due to spatial spill-over or segmentation errors. Since the expression of markers like CD3 can typically be found in a narrow area around the nuclei, while myeloid markers such as CD163 are diffuse and can therefore more easily lead to staining of neighboring cells, we prioritized markers based on their staining. In this specific example we would **label** CD3+CD163+ cells as T cells since we prioritize CD3.
6. **Tissue segmentation:** Cytokeratin (CK) is used as a tumor cell marker. A first “naive” tissue segmentation is done by assigning all cells with CK score above the CK threshold to tumor tissue type and all other cells to stroma tissue type. In a second step, groups of less than 5 stroma (respectively tumor) cells connected to a group of more than 10 tumor (respectively stroma) cells are reassigned to tumor (respectively stroma) tissue.
7. **TLS detection**: As a proxy for TLS, we use patches of B cells (CD20+ cells) with a local B cells density above 2000 cells/mm^2^ and at least 40 cells. Patches are found using the alpha shape^57^ for the set of B cell positions.

A TSV file is generated for each image containing a row for each cell with the summary intensity of each marker, the area of the cell, whether the cell is located in tumor, stroma, or TLS, and its X and Y location.

Reading of the QPTIFF format is performed with the Bio-formats command line tools (https://www.openmicroscopy.org/bio-formats). Image manipulation is performed with libvips (https://libvips.github.io/libvips). The image analysis script is running in the R software (https://www.R-project.org) with the help of multiple libraries. The web-tool is developed in PHP (backend) and in Javascript (Frontend). The backend uses the SLIM framework (https://www.slimframework.com, version 3). Cell quantification data is loaded and indexed in a SQLite3 database for improved performance. The tiled images are served by the IIPImage image server, using the Internet Imaging Protocol. The frontend is developed with the VueJS (https://vuejs.org/, version 2) and the Bootstrap (https://getbootstrap.com/, version 4) libraries. The javascript image viewer is based on the Openseadragon library (https://openseadragon.github.io/, version 4), with the OpenSeadragonFiltering (https://github.com/usnistgov/OpenSeadragonFiltering) and OpenSeadragonScalebar (https://github.com/usnistgov/OpenSeadragonScalebar) plugins. The FabricJS library (http://fabricjs.com/, version 5) is used to draw annotations on the image. The source code of the web-tool as well as the different components are available as a docker image (https://github.com/BICC-UNIL-EPFL/IFQuant).

### IMC data preprocessing

MCD files from IMC were processed using the steinbock toolkit (v0.14.1)^34^. First, TIFF files were generated and hot pixel filtered. Segmentation was performed using the Mesmer^58^ implementation within steinbock. Histone H3 and Iridium were used for nuclei detection and E-cadherin, CD3, CD8, CD20, CD163, were used as markers for cellular boundaries. Cellular interactions were quantified as cells that touch after a cell boundary expansion of 4 µm. Finally, the mean marker intensities, cellular area, centroid, major and minor axis length and eccentricity were calculated per cell. One CSV file per image containing the mean pixel intensities, one CSV per image containing the cellular information and one CSV per image containing the neighboring cell information were exported.

### IMC cytoblock stability analysis

Single cell data was obtained from pre-processing and was analyzed in R. Spillover correction was performed as previously described using the imcRtools and CATALYST packages^59,60^. Graph-based clustering using Phenograph^61^ was performed to identify cell phenotypes using 45 nearest neighbors and clusters were annotated manually. The fractions of cell phenotypes measured in cytoblocks with and without activation were calculated per batch and the coefficients of variation calculated across batches. To compare the marker expression variation across batches, for each cell type we only considered markers expressed in that cell type to avoid calculating variation of noise (**Supplementary Table 3**). The coefficients of variation of each of the cell type relevant markers were calculated per cell phenotype across batches.

### IMC cell phenotype classification

A total of 179 images from 179 patients spanning the five tumor indications were selected for cell phenotype annotation. The R Bioconductor package cytomapper^62^ was used to label cells via multi-dimensional gating and to inspect gates on images. A detailed sketch of the gating strategy can be found in **Supplementary Methods 2**. Slightly over 2000 gates were stored for 14 cell phenotypes and a total of 340’000 cells.

All gates were then concatenated and cells that had been labelled twice were excluded if they were not of type tumor. Cells that were labeled as tumor and another cell phenotype were finally labelled as the other cell phenotype to preserve immune cells within the tumor compartment which may be tumor marker positive due to spatial marker spillover. A random forest classifier was trained using the caret^63^ package with the cancer indication as a covariate. Briefly, the data was split into training (80%) and test data set (20%) and a five-fold cross validation was used to train the model while tuning the mtry parameter. The model was then tested on the test data set (**Supplementary Figure 4A**). The pre-trained random forest classifier was applied to all samples stained with panel 1 in IMMUcan and the 10 samples presented here. The labelled data used for training of the classifier can be found *ZENODO_link*.

### IMC data analysis and comparison with IF

IMC data analysis was performed using R. The imcRtools^34^ package was used to read pre-processed data from steinbock following our workflow^34^. CD20 patches were calculated based on the neighborhood graph calculated with steinbock and patches were called when consisting of minimally 25 B or BnT cells. Detected patches were expanded by 10µm to include closely neighboring cells. Tumor patches were also calculated based on the neighborhood graph from steinbock and had to initially consist of 25 tumor cells. We also consider all cells within 25 µm from a tumor patch cell to belong to tumor patches. For comparison with the mIF data, the ROIs from IMC were manually identified in the mIF images using the napari *affinder* plugin (https://www.napari-hub.org/plugins/affinder). mIF and IMC images were then aligned using a similarity transform with the landmark points obtained from affinder. Of note, one image was removed due to imprecise alignment. Matched cell phenotypes between both datasets were used for direct comparison. Differential abundances of cell phenotype counts between mIF and IMC images were calculated using the edgeR^64^ Bioconductor package. Spatial associations between matched cell phenotypes in mIF and IMC from the consecutive sections were calculated using the L-cross function from the sf and spatstat^65^ R packages. We performed a maximum absolute deviation (MAD) test to assess if the estimated L function wanders outside an envelope around the expected L function generated by calculating the L function of 99 simulated realizations of complete spatial randomness. We consider cell phenotype pairs per image set to be similarly co-localised between mIF and IMC if the area between the estimated and expected L function is larger than 0, and if the MAD test p-value is smaller than or equal to 0.01. We used the spicyR^38^ Bioconductor package to detect differential cell phentopye/cell phenotype co-localization between IMC and mIF at a false discovery rate threshold of 10%.

## Data and Software availability

● Upload mIF and IMC data to Zenodo
● Upload classifier to zenodo
● Links to code on github
● Links to containers of docker hub

