## supplementary material for "Multi-modal image analysis for large scale cancer tissue studies within IMMUcan"

Supplements

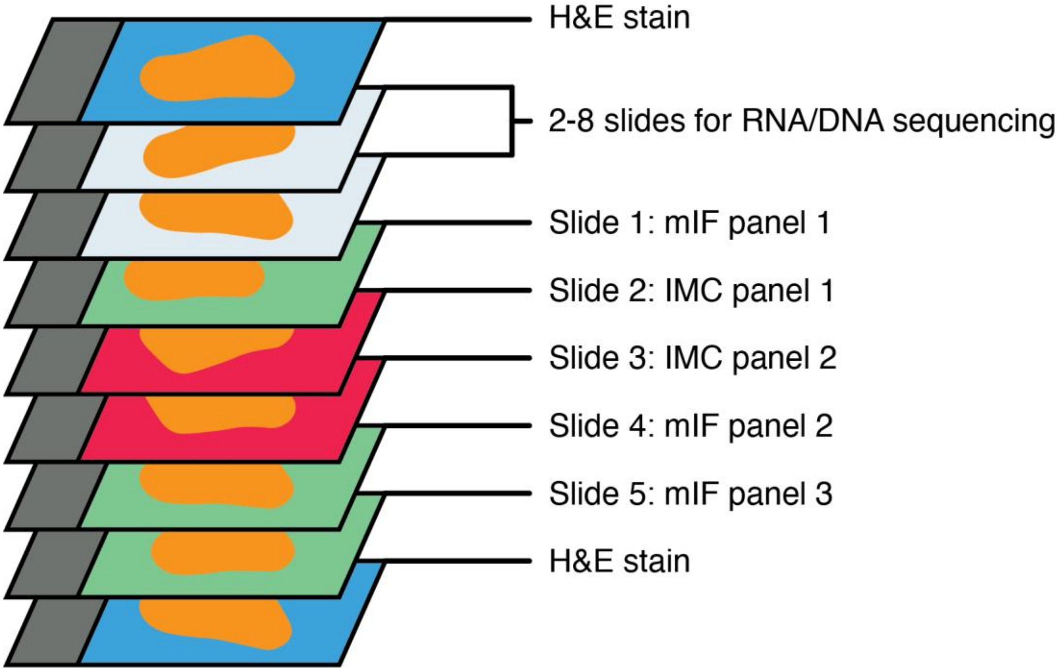


**Supplementary Figure 1: FFPE processing for IMMUcan**.

FFPE blocks from eligible patients were collected and sectioned as shown above. Depending on the size of the tumor a variable number of sections were cut for RNAseq and WES.

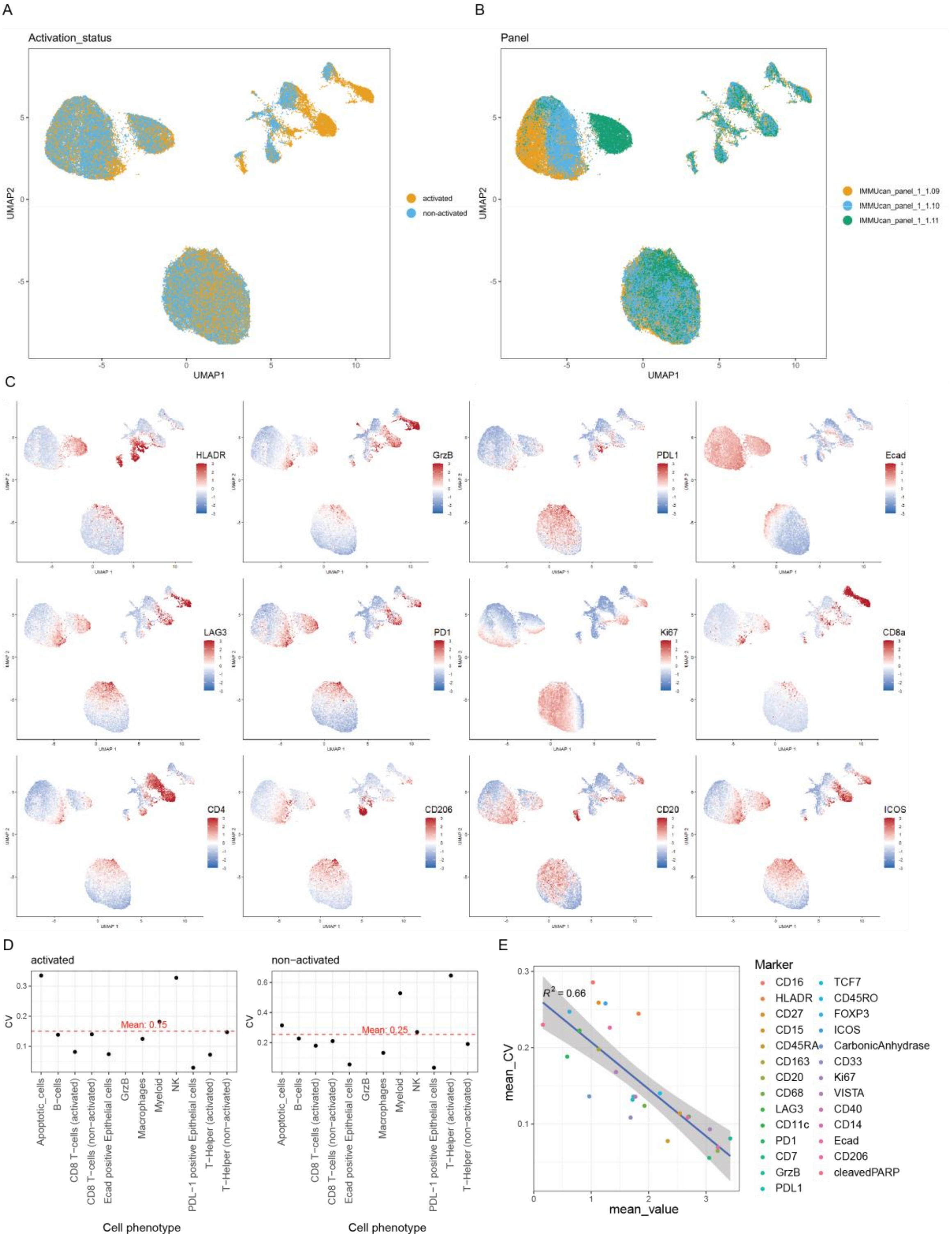


**Supplementary Figure 2: Cytoblock based IMC measurement stability**.

**A.** UMAP of single cells from cytoblocks colored by the activation of the PBMCs in the cytoblocks. **B.** UMAP as in A colored by the antibody panel mix used for staining. **C.** UMAP of single cells from cytoblocks colored by the z-scaled expression of markers. **D.** For each cell phenotype detected in the cytoblocks the mean coefficient of variation (CV) of the detection frequency over all time points (sample batches) is shown (black points) for activated cytoblocks on the left and for non-activated cytoblocks on the right. The Horizontal red dashed lines indicate the mean observed CV for all cell phenotypes. **E.** scatterplot of the observed mean expression of each marker on the x-axis and the average coefficient of varation for each marker calculated across batches on the y-axis. Individual points are colored by the respective marker.

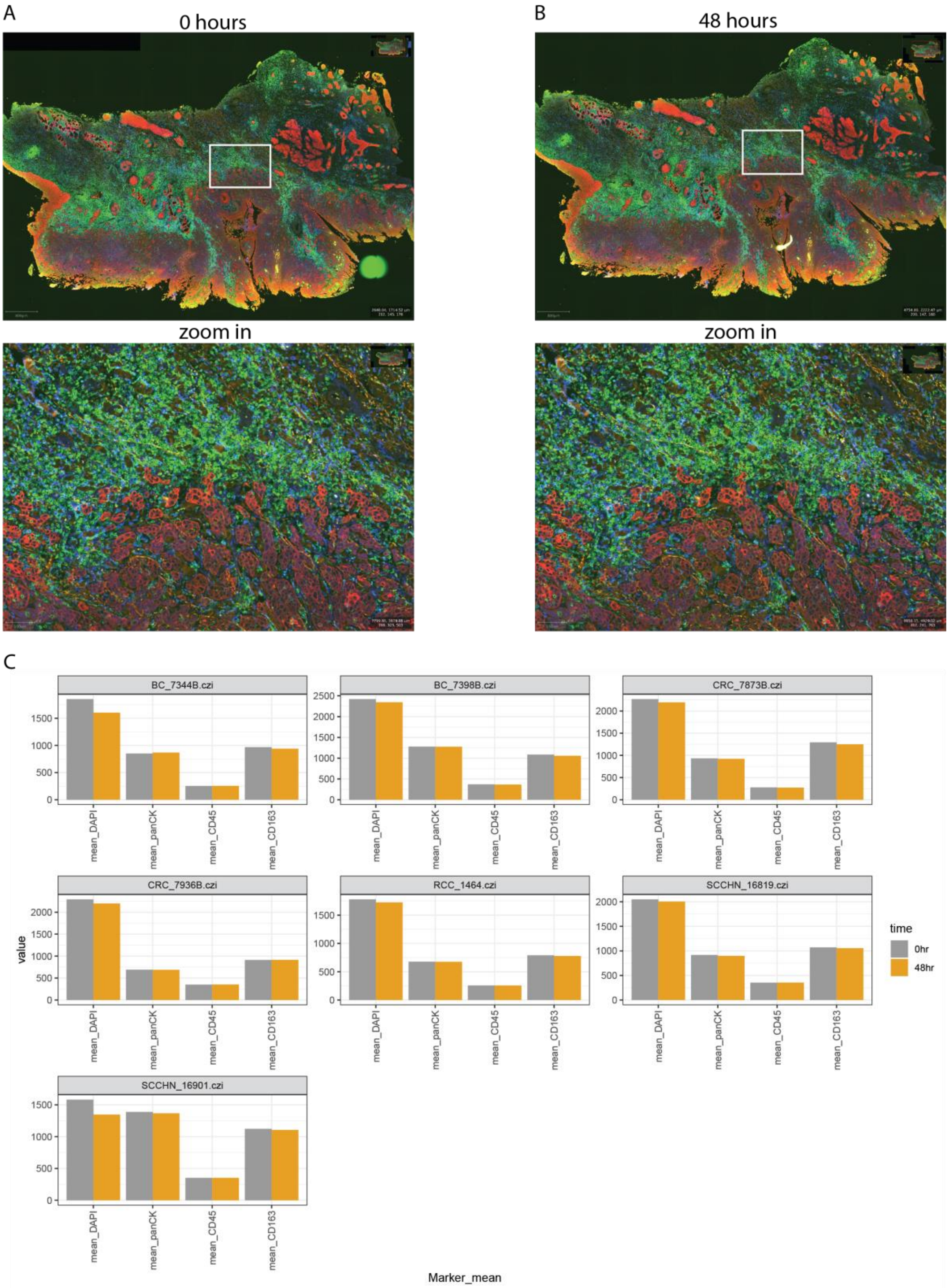


**Supplementary Figure 3: Fluorescence signal stability in dry format**.

Three color fluorescence images recorded immediately after drying (**A**) and 48 hours after drying (**B**). Zoom ins are shown on the bottom. Scale bars on the top row are 800 µm and on the bottom row 100 µm. **C.** The fluorescence intensity in segmented single cells for 7 individual tumors at 0 and 48 hours after drying for Dapi, pan Cytokeratin, CD45 and CD163 is shown.

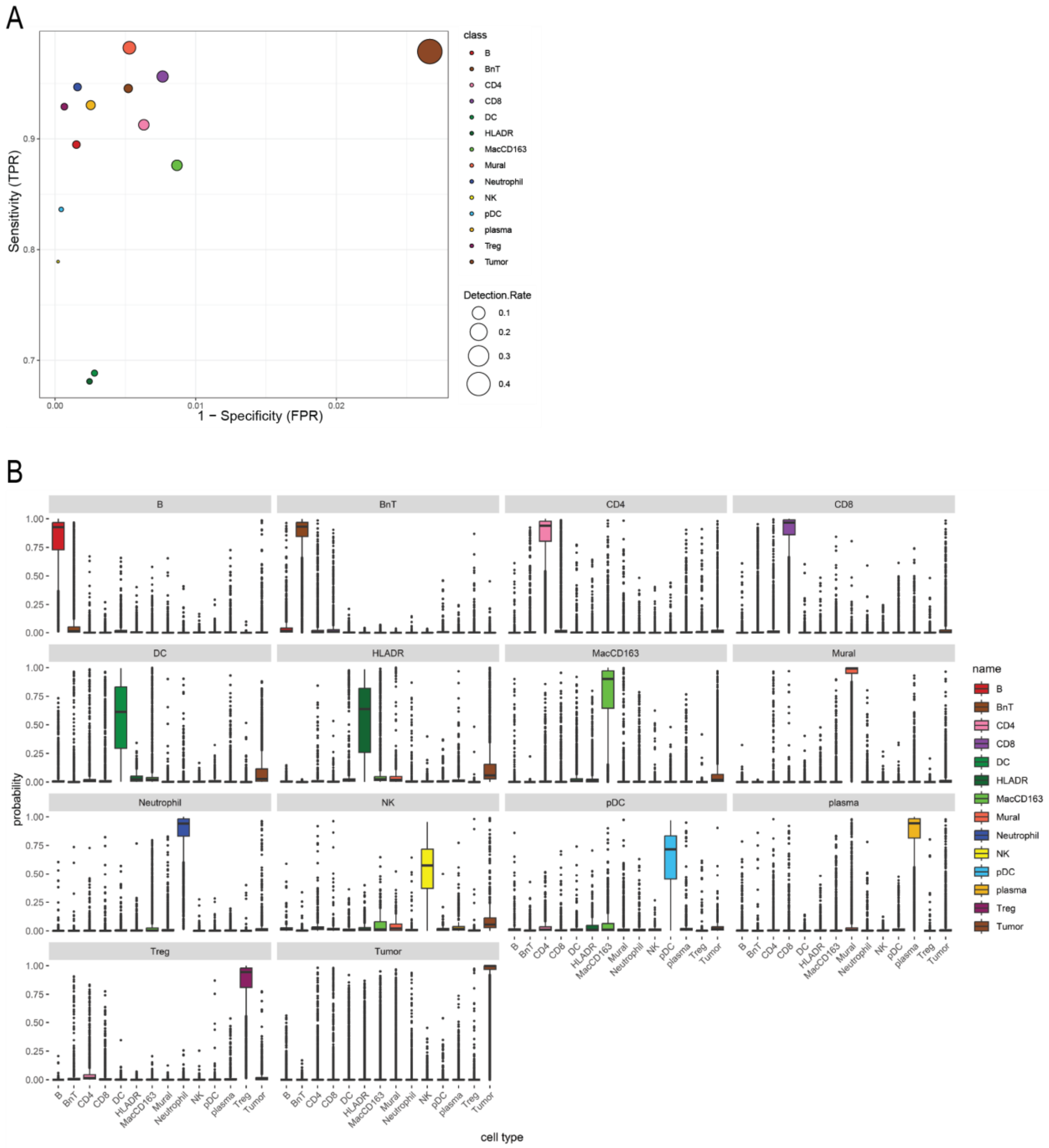


**Supplementary Figure 4: Random forest classifier for cell types.**

**A.** True positive rates (TPR) and false positive rates (FPR) for the detection of individual cell types are shown based on a hold out test data set. **B.** For each ground truth labelled cell type the probabilites of all cells of this type belonging to any of the classes on the x-axis are shown.

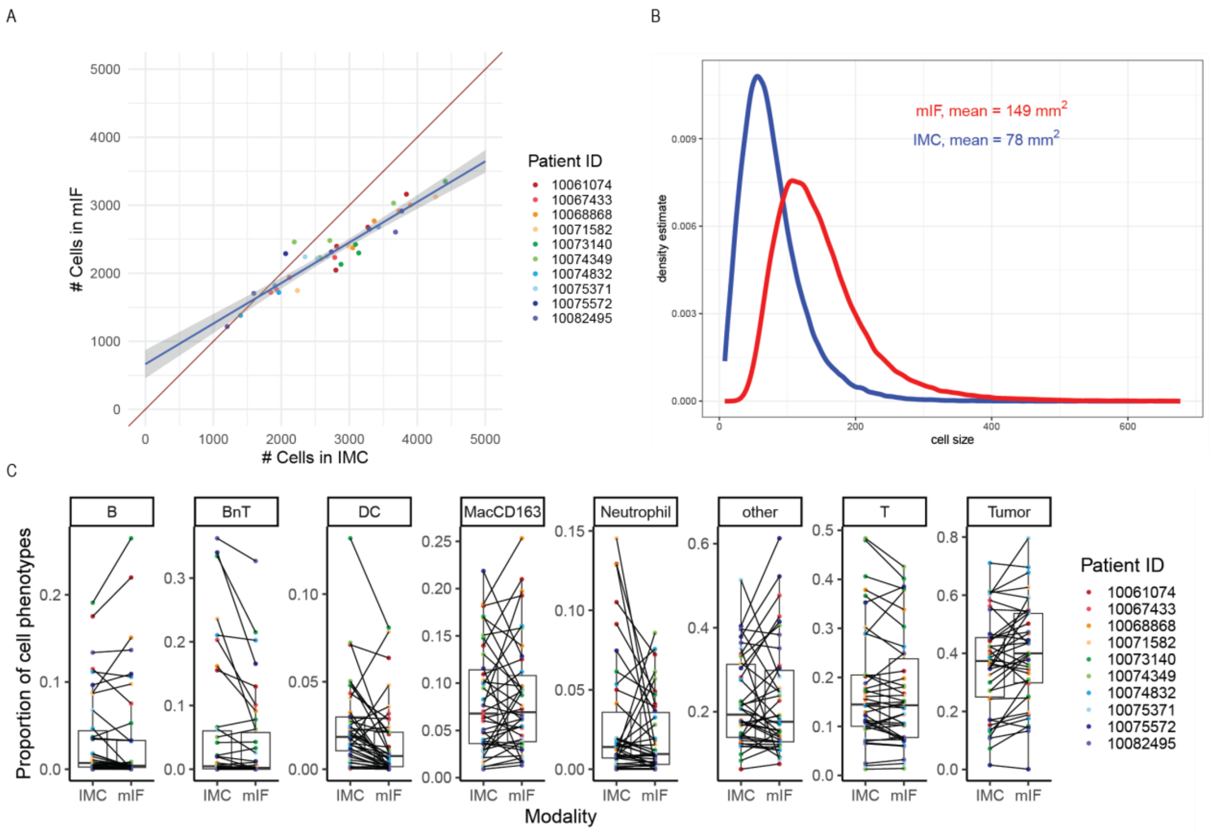


**Supplementary Figure 5: Comparison of IF and IMC.**

**A.** Scatterplot showing the number of segmented cells in IMC on the x-axis and in mIF of the matched region on the y-axis. Images are colored by patient ID. **B.** Distribution of the single cell area in mIF (red) and IMC (blue). **C.** For each matched cell phenotype the proportion of the phenotype among all cell phenotypes is compared between mIF and IMC for matched images.

A

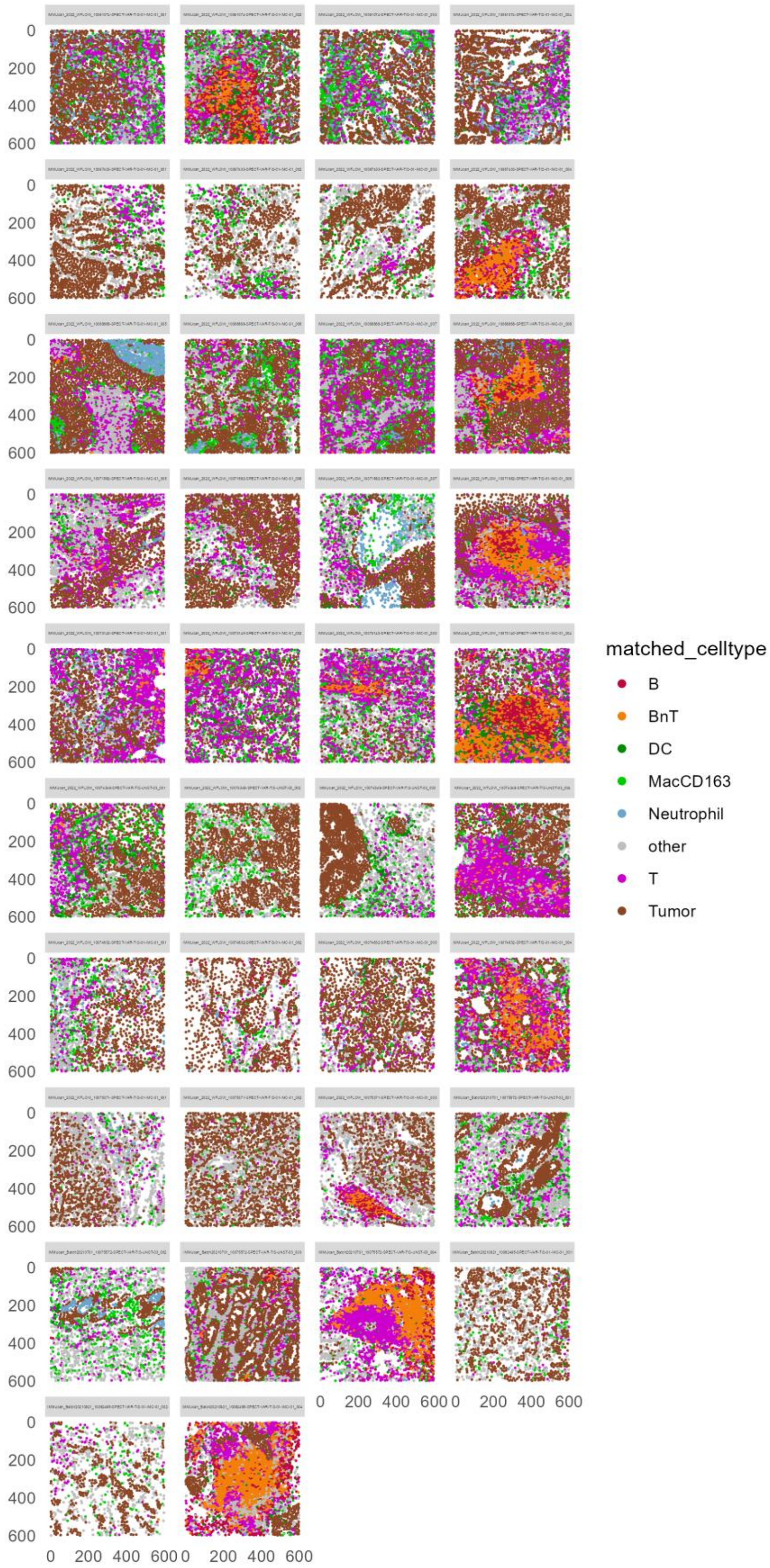
matched_celltype
B
BnT
DC
MacCD163
Neutrophil
other
T
Tumor
0
200
400
600

B

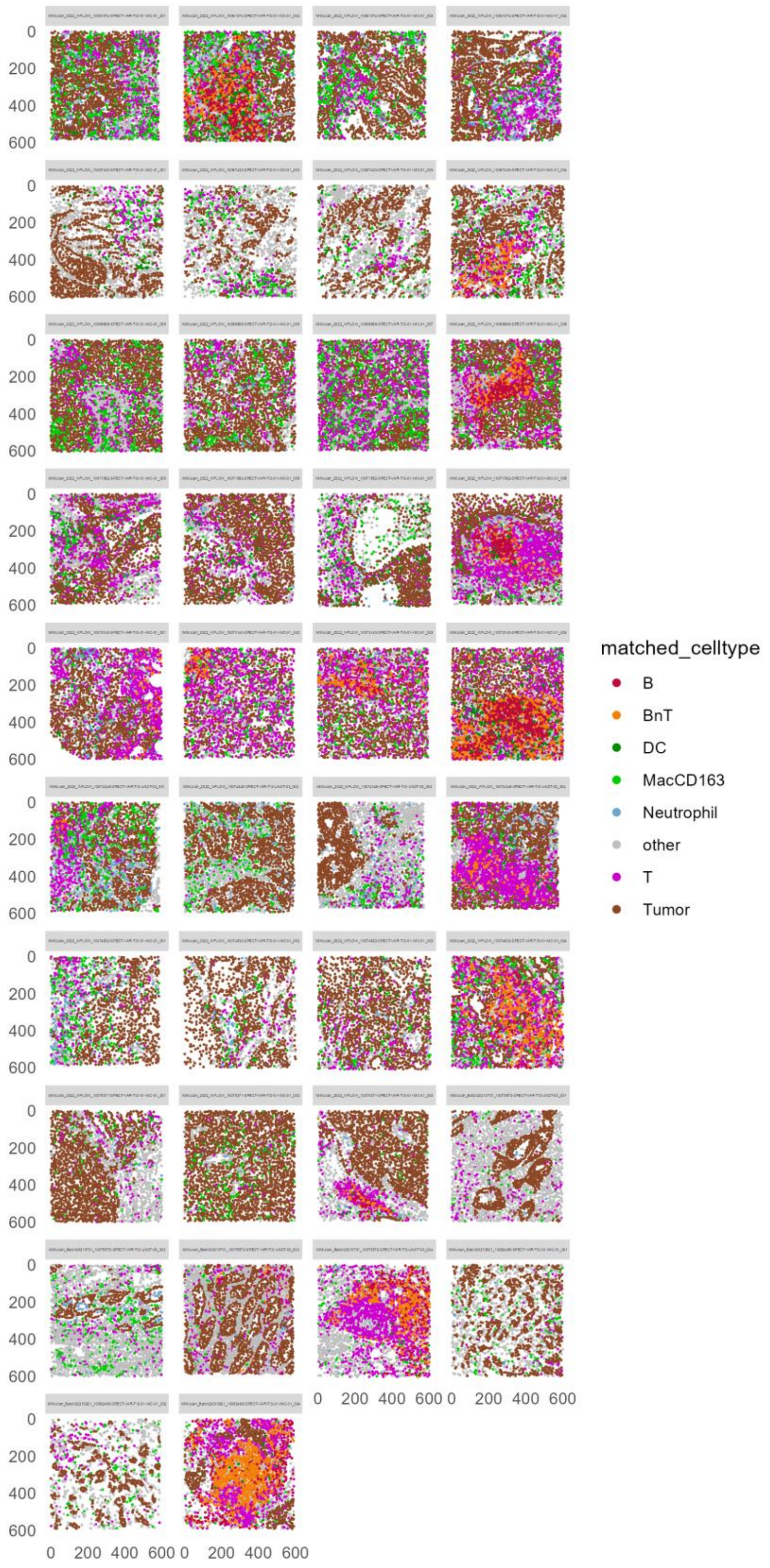


**Supplementary Figure 6: Example images of segmented single cells from mIF and IMC.**

**A.** Examples of single cells on images. Each cell is represented by a point reflecting the center of the cell and colored by the respective cell phenotype. Dimensions of the images are given on the x and y axis. **B.** As in A for mIF.

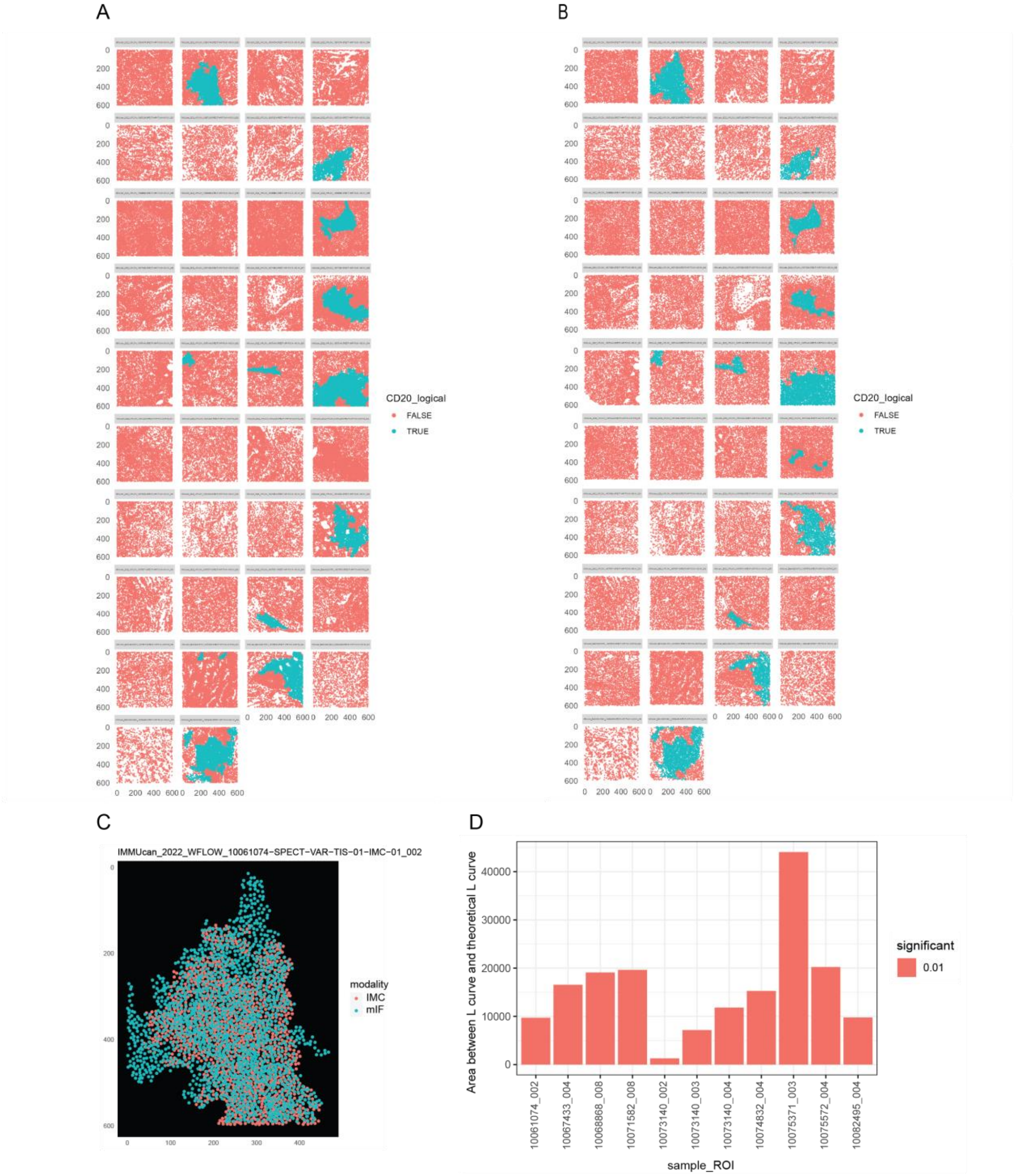


**Supplementary Figure 7: spatial comparison of mIF and IMC.**

**A.** B cell patch detection in IMC based on CD20 positive cells. Cells are depicted as points by the center **B.** B cell patch detection in mIF based on CD20 positive cells. **C.** Center points from B cells of one matched image from mIF (green points) and IMC (red points) were overlayed to highlight the matched location of the cells in the image. The image corresponds to the image from the top row, second column in A and B. **D.** Results from Lcross functions applied to B cells for mIF and IMC are shown as the area between the calculated and the theoretical L curve. Positive values indicate that cells are more clustered as expected if cells were randomly distributed in the images. Significances were calculated based on 100 envelope simulations.

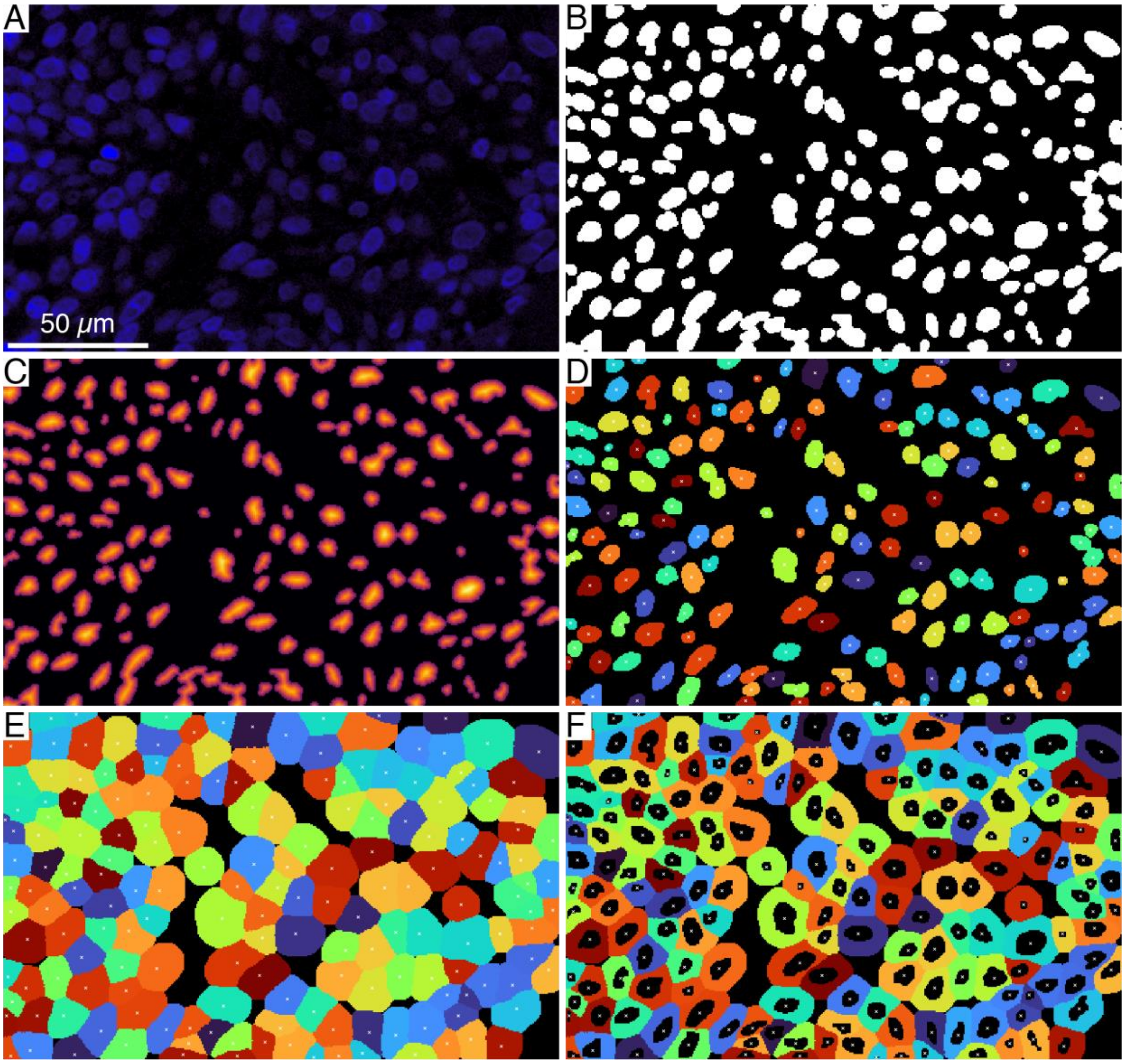


**Supplementary Figure 8: Nuclei & cell segmentation.**

**A.** DAPI channel. **B.** Nuclear mask obtained by adaptive thresholding. This mask contains value 1 (white) for nucleus regions and 0 (black) for background. **C.** Distance map of the nuclear mask colored from black (distance 0) to yellow (maximum distance). **D.** Nuclear mask after cleaning (colored by cell ID, background in black) with nuclei centers (white crosses). **E.** Cell regions (colored by cell ID, background in black) approximated by Voronoi based segmentation with nuclei centers (white crosses). **F.** Cytoplasm regions (colored by cell ID, background in black) with nuclei centers (white crosses).

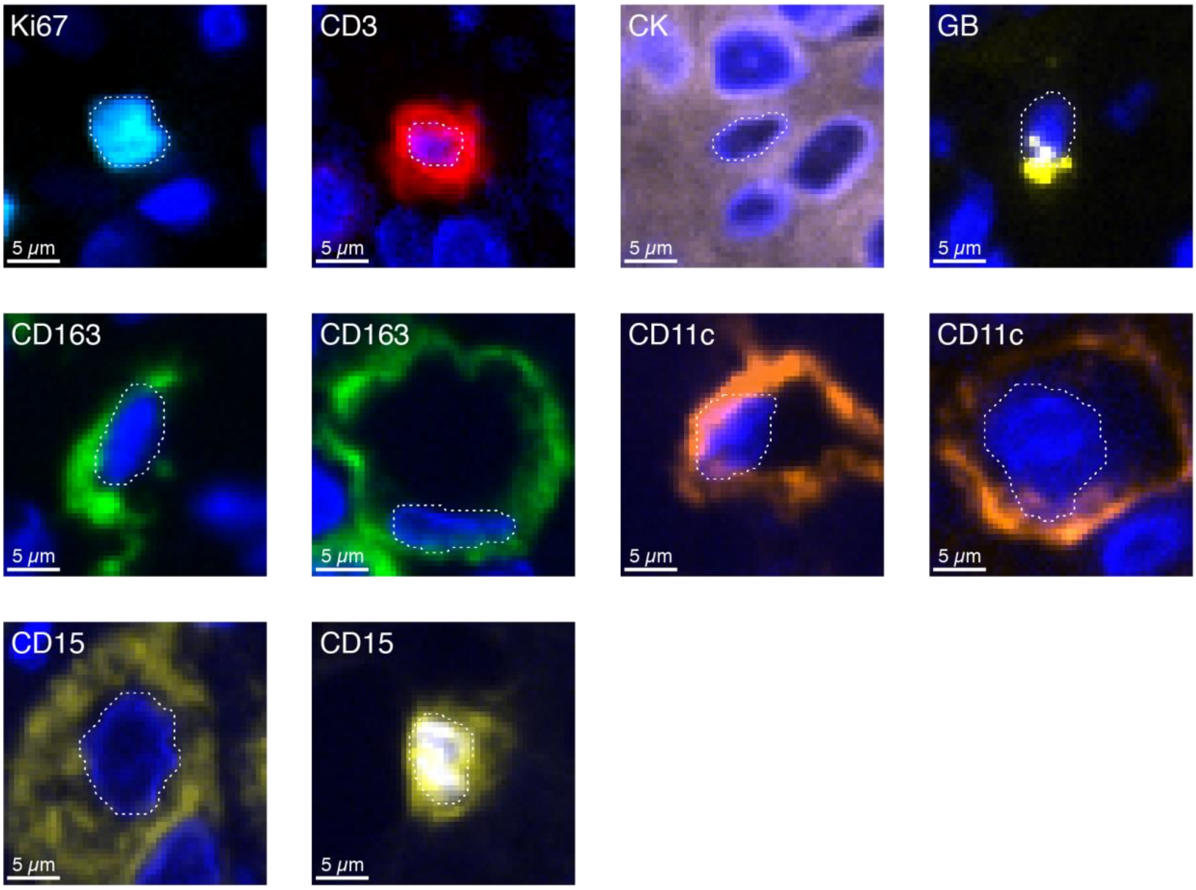


**Supplementary Figure 9: Spatial distribution of fluorescence for various markers.**

Each panel shows DAPI channel (blue) with nucleus border (dashed white) together with the fluorescence for specific marker (label in upper-left corner): Ki67 (cyan), CD3 (red), CK (pink), GB (yellow), CD163 (green), CD11c (orange) and CD15 (yellow).

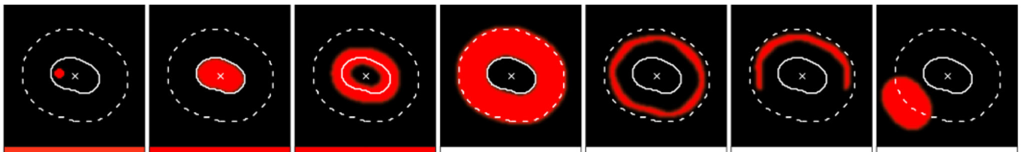

| | | | | | | | |
|---|---|---|---|---|---|---|---|
| 95–th percentile (nucleus) | 0.9 | 1 | 1 | 0 | 0 | 0 | 0 |
| 75–th percentile (nucleus) | 0 | 1 | 1 | 0 | 0 | 0 | 0 |
| median (nucleus) | 0 | 1 | 1 | 0 | 0 | 0 | 0 |
| Encircling median (nucleus) | 0 | 1 | 1 | 0 | 0 | 0 | 0 |
| 95–th percentile (cytoplasm) | 0 | 0 | 1 | 1 | 1 | 0.9 | 1 |
| 75–th percentile (cytoplasm) | 0 | 0 | 0.7 | 1 | 0.6 | 0.07 | 0 |
| median (cytoplasm) | 0 | 0 | 0 | 1 | 0.008 | 0 | 0 |
| Encircling median (cytoplasm) | 0 | 0 | 1 | 1 | 1 | 1 | 0 |
| 95–th percentile (cell) | 0 | 1 | 1 | 1 | 1 | 0.9 | 1 |
| 75–th percentile (cell) | 0 | 0 | 0.9 | 1 | 0.3 | 0.04 | 0 |
| median (cell) | 0 | 0 | 0 | 1 | 0.001 | 0 | 0 |
| Encircling median (cell) | 0 | 1 | 1 | 1 | 1 | 1 | 0 |

**Supplementary Figure 10: Single cell quantifications in mIF.**

Score obtained with various combinations of summary statistics and regions (rows) evaluated on archetypal spatial distributions of fluorescence (columns). Images on top show distributions of fluorescence, colored from black (fluorescence=0) to red (maximum fluorescence=1), with nucleus center (white cross), nuclear region boundary (plain lines) and cell region boundary (dashed line).

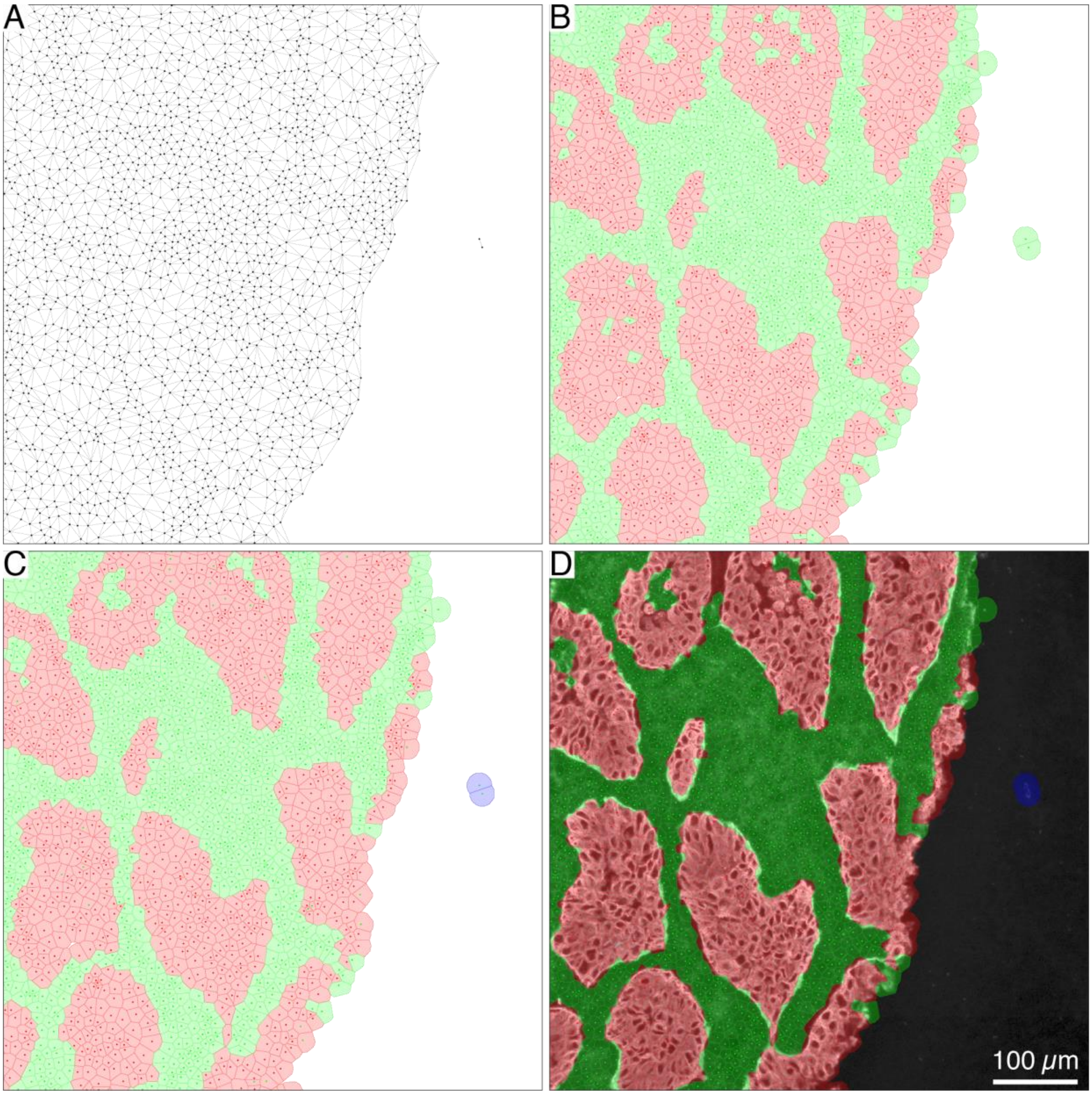


**Supplementary Figure 11: Tissue segmentation.**

**A.** Delaunay triangulation. Edges with length above 40 μm are not shown. Vertices (black dots) correspond to nuclei centers. **B.** Voronoi tessellation after clipping each cell to a maximum distance to nucleus center of 15 μm. Nucleus centers (dots) and Voronoi cells (polygons) are colored red for CK positive cells and green for CK negative cells. **C.** Clipped Voronoi tessellation with nucleus centers colored by CK status (red for CK positive cells, green for CK negative cells) and Voronoi cells colored by final tissue type (stroma in green, tumor in red and "other" in blue). **D.** CK channel colored from black (no CK) to white (maximum CK) with clipped Voronoi tessellation from C overlaid.

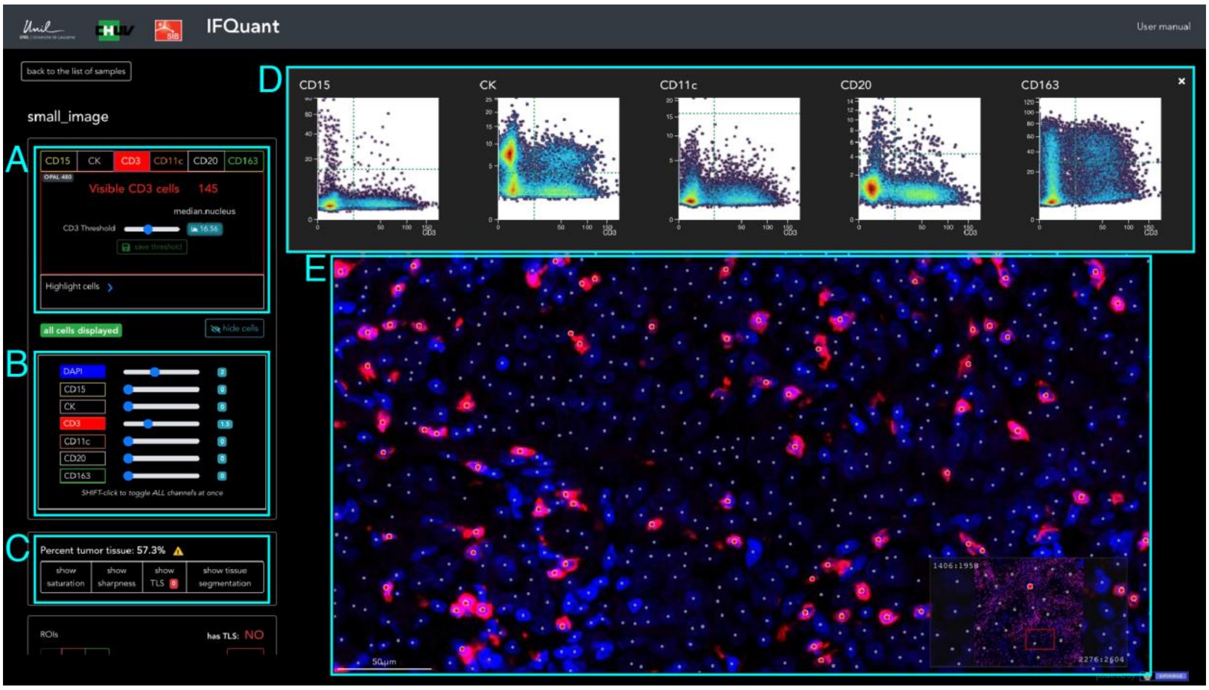


**Supplementary Figure 12: Web application.**

**A.** List of markers. Threshold review/adjustment is done one marker after another. **B.** Enable to combine channels in a composite image. **C.** Display QC, TLS and tissue segmentation masks. **D.** Scatter plots of marker scores for all markers (y-axis) versus the selected marker (x-axis). Marker thresholds are depicted in dotted lines. **E.** Composite image. Positive cells for the selected marker are flagged with a red circle. Negative cells with a smaller grey circle.
